# Central Complex representations of self-movement are sufficient to compute wind direction in flight

**DOI:** 10.1101/2025.05.09.653128

**Authors:** Christina E. May, Benjamin Cellini, S. David Stupski, Austin Lopez, Nehal Mangat, Floris van Breugel, Katherine I. Nagel

## Abstract

External forces such as gravity and wind fundamentally shape how an animal navigates through the world, but force magnitude and direction cannot be directly measured by an animal in motion. How the brain integrates multiple sensory cues to determine the direction of such external forces remains unclear. In the fruit fly (*Drosophila melanogaster*) columnar inputs to the navigation center, known as PFNs, have been shown to represent both visual and mechano-sensory cues about self-motion in a similar vector format, but the dynamics and integration of these cues, particularly during complex natural movements, are not known. Here we used 2-photon imaging to investigate the dynamic representation and integration of multi-sensory self-motion cues – optic flow and airflow – across PFNs. We find that one type, called PFNd, linearly integrates optic flow and airflow direction signals with distinct dynamics. We next reveal airflow speed as well as direction encoding by PFNp_c neurons. Imaging from available genetic lines suggests that many PFN types likely encode functions of airflow speed and direction. Based on these data, we construct and validate computational models of how visual and mechano-sensory dynamics are integrated and represented across PFN populations. We then use these models to estimate neural responses during real and simulated flight maneuvers. Applying a nonlinear empirical observability analysis to these responses, we show that active turning and deceleration maneuvers combined with PFN representations can decode the direction of an ambient external cue (wind) during free flight, and identify the theoretical contribution of different PFNs to this ability. Finally, we show that an anatomically-inspired artificial neural network can successfully decode wind direction from PFN representations and identify components of the algorithm used by both the network and real flies to orient upwind during olfactory navigation in flight. Our work provides evidence that active sensation, combined with multi-sensory encoding, can allow a compact system like the fly navigation center to infer a property of the external world that cannot be directly measured by a single sensory system.

## Introduction

External forces such as gravity and wind serve as crucial orientation cues for navigation. For example, flying insects use wind direction to navigate towards odor sources both while walking and in flight (*1–5*). Inferring the direction of such a force is challenging when an animal is in motion, because these forces cannot be directly detected by an animal’s sensors. Instead, external forces sum with self-generated motion to produce patterns of sensory feedback in both visual and mechano-sensory modalities (*6*, *7*). By combining active maneuvers with multi-sensory feedback, recent theoretical studies have argued that an animal should be able to infer the direction of wind while in flight (*8–11*). Whether neural representations of self-motion are sufficient for such calculations, and what computations transform sensory feedback into estimates of such external world variables are unknown.

When an animal moves through the world, it produces both characteristic patterns of visual feedback, known as optic flow (*12*, *13*), and patterns of mechano-sensory feedback, in the form of vestibular sensation (*14*). Optic flow provides high directional accuracy but is relatively slow, can be evoked artificially by environmental movement, and has a complex relationship to ground speed (*8*, *15*, *16*). In contrast, vestibular feedback is fast, but only transiently encodes movement speed, acceleration, and direction (*17*). While both cues can redundantly signal self-motion in some cases, neither cue alone is sufficient to accurately estimate self-motion under natural conditions (*16*).

Consistent with their prominent role in self-motion estimation, neurons that integrate vestibular and visual self-motion cues have been described in many regions of the vertebrate brain, including the vestibular brainstem (*16*, *17*), cerebellum (*18–20*), and temporal and parietal cortices (*21–23*). How these multimodal representations are translated into internal estimates of navigational variables remains challenging to address in vertebrate brains.

In insects, columnar neurons called PFNs that input to the brain’s navigation center (Central Complex, CX) have been shown to encode both optic flow direction (*24–26*) and airflow direction (*27*, *28*) – visual and mechano-sensory cues that both signal self-motion (*8*, *29*, *30*). Intriguingly, both modalities appear to be represented in a similar “basis vector” format (Fig. 1A) (*24*, *27*, *31*, *32*). In this format, PFNs of the same type on each side of the brain represent basis vectors oriented at +/- 45° to the fly’s heading (Fig 1A: dotted blue and red lines). The fly’s heading relative to the world is represented by the location of a bump of Ca^2+^ activity across each array of neurons (Fig 1A: location of sinusoid peak and shaded circles below). The direction of optic flow or airflow (Fig 1A: solid blue and red lines) is projected onto these two basis vectors to produce different bump amplitudes in the two hemispheres (Fig 1A: amplitudes of sinusoids). Thus, the location and relative amplitude of activity in each PFN type represents the airflow or optic flow direction relative to the fly. Through summation of specific sets of PFN types, recent studies have shown that the CX can perform vector addition to compute the fly’s allocentric traveling direction (*24*, *31*), or theoretically compute the allocentric direction of the wind while walking (*28*). Whether this type of vector representation is found across PFN types, and how visual and mechano-sensory cues are represented and integrated during dynamic input are not known.

**Figure 1.**
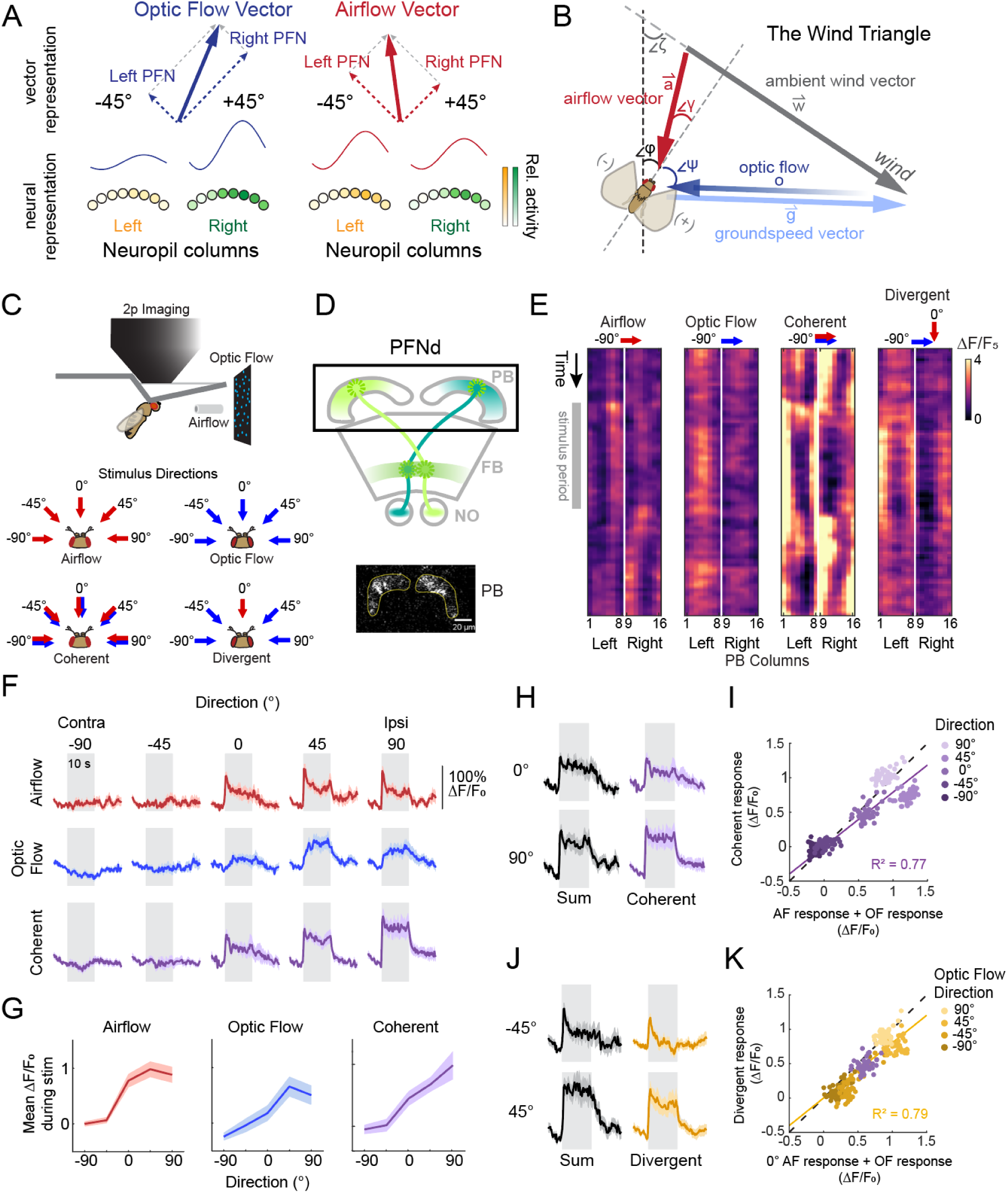
PFNd neurons integrate optic flow and airflow direction with distinct timecourses. A. Vector representations by calcium activity across populations of columnar neurons (PFNs). Bump position represents allocentric heading angle while the relative amplitude of left and right bumps represents egocentric direction of optic flow or airflow (*24*, *31*, *32*). B. The wind triangle for flying flies. Ambient wind produces offset airflow (*red*) and optic flow (*blue*) vectors for a flying fly. Ambient wind direction (ζ) in allocentric coordinates and wind speed (w) determine the egocentric airflow direction (γ) and airspeed (a), and egocentric travel direction (ψ) and groundspeed (g). The optic flow vector indicates the direction and magnitude for ventral optic flow over flat terrain, and points in the opposite direction from the travel direction, and its magnitude (o) is equal to g divided by altitude. C. *(above*) Paradigm for two-photon imaging of responses to directional airflow and optic flow in tethered, non-flying flies. (*below*) Stimuli of each modality were presented from five directions alone, coherently, or from diverging directions. D. PFNd anatomy. (*above*) Anatomy of PFN neurons: these receive input in the protocerebral bridge (PB) and noduli (NO) and output in the fan-shaped body (FB). Bump-shaped calcium activity can be imaged in the PB or FB (green and blue gradients). Imaging was performed from the PB as indicated by the black box. (*below*) Single volumetric (mean z-projection) frame of imaging from the PB in PFNd showing bump structures in left and right hemispheres. ROIs of the left and right PB halves indicated in yellow. E. Example single-trial calcium activity in PFNd PB hemispheres in response to select stimuli. All data from the same fly. F. Calcium activity dynamics in PFNd PB halves to contralateral (negative directions) and ipsilateral (positive directions) stimuli. Gray areas indicate 10-s stimulus period. Colored traces represent mean response ±SEM across flies (N=10 flies, n = 2-4 trials per fly). G. Mean calcium response in PFNd PB halves as a function of stimulus direction for airflow only (*left*: 1^st^ second of response), optic flow only (*middle*: last 3 s of response), and coherent multimodal stimuli (*right*: full 10-s stimulus period). Positive values represent ipsilateral stimuli and negative values represent contralateral stimuli as in F. Traces represent mean ±SEM. Same data as in F. H. Comparison of the sum of responses to individual stimuli from the same direction (black traces, left panels) to responses to coherent stimuli (purple traces, right panels). Traces represent mean ±SEM across flies (N=10 flies, n= 2-4 trials), same data as F. I. The sum of the 10-second mean responses to individual stimuli (x-axis) versus the mean activity during the 10-second coherent stimulus (y-axis). Each dot is one timepoint of the mean across flies for each stimulus direction. Regression line in purple. N=10 flies, same data as F. J. Comparison of the sum of responses to individual stimuli from different directions (black traces, left panels, OF direction indicated and AF direction = 0°) to responses to divergent stimuli (purple traces, right panels). Traces represent mean ±SEM across flies (N=10 flies, n= 2-4 trials), same data as F. K. The sum of the 10-s responses to divergent individual stimuli (x-axis) versus the mean activity during the 10-second divergent stimulus (y-axis). Each dot is one timepoint of the mean across flies for each stimulus direction. Regression line in yellow. n=10 flies, same data as F.

One computation that might take advantage of this multi-sensory representation is resolving the direction of ambient wind during flight. During flight, wind direction, groundspeed, airflow, and heading are related through the so-called wind triangle (Fig. 1B). If sufficient angles or lengths of this triangle are known, the ambient wind direction can be decoded accurately (*8*, *11*). For example, if one knows both the angles of egocentric self-motion (*ψ*) and airflow (*γ*), and the lengths of the airflow (*a*) and groundspeed (*g*) vectors, the ambient wind direction can be estimated through vector subtraction. However, the true groundspeed vector is challenging to estimate, because its relationship to optic flow speed depends on the distance of visual objects. Without the length of this vector, the ambient wind direction is non-trivial to calculate. An alternative approach is to make two or more measurements, separated by a change in speed or direction. If the fly changes speed or direction, effectively changing the airflow and groundspeed vectors while the wind vector remains the same, it could theoretically derive the ambient wind direction from only partial measurements of the triangle (*11*, *33*). Behavioral studies support the idea that flying flies estimate wind direction by combining sensory feedback with active maneuvers (*33*): flies whose olfactory receptor neurons are activated opto-genetically make rapid (100-ms) and ballistic turns upwind. The speed of this movement argues that it cannot depend on slow visual feedback (*34*, *35*) but more likely uses an internal estimate of the ambient wind direction. Upwind turns are often preceded by either a deceleration or a turn in a random direction (*33*). These maneuvers could provide the second measurement that allows the fly to deduce the ambient wind direction.

We hypothesized that the multi-sensory representation of self-motion generated in PFNs could provide a computational substrate for estimating wind direction while in flight. To test the feasibility of this hypothesis, we used 2-photon imaging to measure the dynamic integration of optic flow and airflow cues across PFNs. We show that one PFN type, called PFNd, linearly sums a transient airflow direction signal with a slower optic flow direction signal, while a second PFN type, called PFNp_c, encodes both airflow direction and speed. We identify a range of different encodings of airflow variables across other PFN types, and show that PFNs remain active during tethered flight, but do not encode turning maneuvers in this setting. Based on our imaging data, we construct dynamic encoding models for PFNs and use these to simulate neural responses to real flight trajectories. We then use a nonlinear empirical observability analysis to show that this representation— when combined with realistic flight maneuvers— is sufficient to allow a fly to decode wind direction. Inspired by the connectivity of PFNs within the CX, we construct a simple feedforward artificial neural network, and show that this network can estimate wind direction from PFN encoding patterns during simulated flight maneuvers. Finally, by analyzing the inputs to our model, we identify two solution regimes to the problem of estimating wind in flight and show that both the network and real flies exhibit differences in performance across these two regimes. Together our data and analysis show how active movements can be combined with multi-sensory encoding to infer the presence and direction of an external force, and provide testable hypotheses about the role of different neuron types in this computation.

## Results

### Approximately linear summation of airflow and optic flow direction signals in PFNd

To examine the dynamic integration of optic flow and airflow cues in PFNs, we presented translational optic flow, airflow, and coherent or divergent combinations of these stimuli to tethered, non-flying flies (Fig. 1C). PFNs (Fig. 1D) make outputs in the Fan-shaped Body (FB) and receive inputs in both the Protocerebral Bridge (PB), and the Noduli (NO). We imaged primarily from the PB (Fig. 1D, lower), in which a bump of activity can be observed in each hemisphere (*24*, *31*). In PFNd neurons, these bumps have been shown to track heading when flies walk or fly in closed-loop with a visual landmark cue, using signals inherited from EPG compass neurons (*24*, *31*, *32*). PFNd neurons have previously been shown to encode both optic flow direction and airflow direction through the relative amplitude of activity in each hemisphere (*24*, *27*), although the integration of these stimuli has not been investigated.

We observed that in non-flying flies, both optic flow and airflow modulated PFNd signal amplitude, but with distinct time courses (Fig. 1E-G). Airflow drove a rapid increase in signal amplitude in the ipsilateral hemisphere that decayed to a lower steady state within a few seconds. Optic flow also drove an increase in amplitude in the ipsilateral hemisphere, but this response was slower and more sustained. When airflow and optic flow were presented from the same direction, we observed a large response with a roughly square amplitude profile, suggesting that airflow and optic flow responses sum. When airflow was presented from the front with optic flow from different directions, we observed the characteristic transient response to frontal airflow, with an additional sustained component during ipsilateral optic flow delivery, also consistent with summation (Supp Fig. 1A). In addition to its effects on signal amplitude, airflow also influenced bump position in a subset of flies (Supp Fig. 1B-E). We did not observe this effect with optic flow (Supp Fig. 1B-E). This movement might reflect the bump activity of EPGs which have been shown to shift with wind direction in non-behaving flies (*36*).

To test the hypothesis of linear summation across modalities, we plotted mean responses to coherent stimuli (Fig. 1H,I) or to divergent stimuli (Fig. 1,J,K) as a function of the sum of the mean individual modality responses. We found that these data were well-fit by a line (R^2^=0.77 for convergent stimuli and R^2^=0.79 for divergent stimuli), although the data for certain directions (45°) was slightly less than predicted by strict linear summation. The most likely source of these discrepancies is a greater weight on airflow input than optic flow input (Supp Fig. 1F,G), and an additional slow decay that is visible with longer stimuli (see Supp Fig. 4J). We observed weak to no tuning for either airflow or optic flow speed in PFNd neurons (Supp. Fig. 1H,I). We conclude that PFNd encodes a multi-sensory representation of self-motion direction by integrating airflow and optic flow signals tuned to the same direction (∼45° ipsilateral) in an approximately linear fashion.

**Figure 2.**
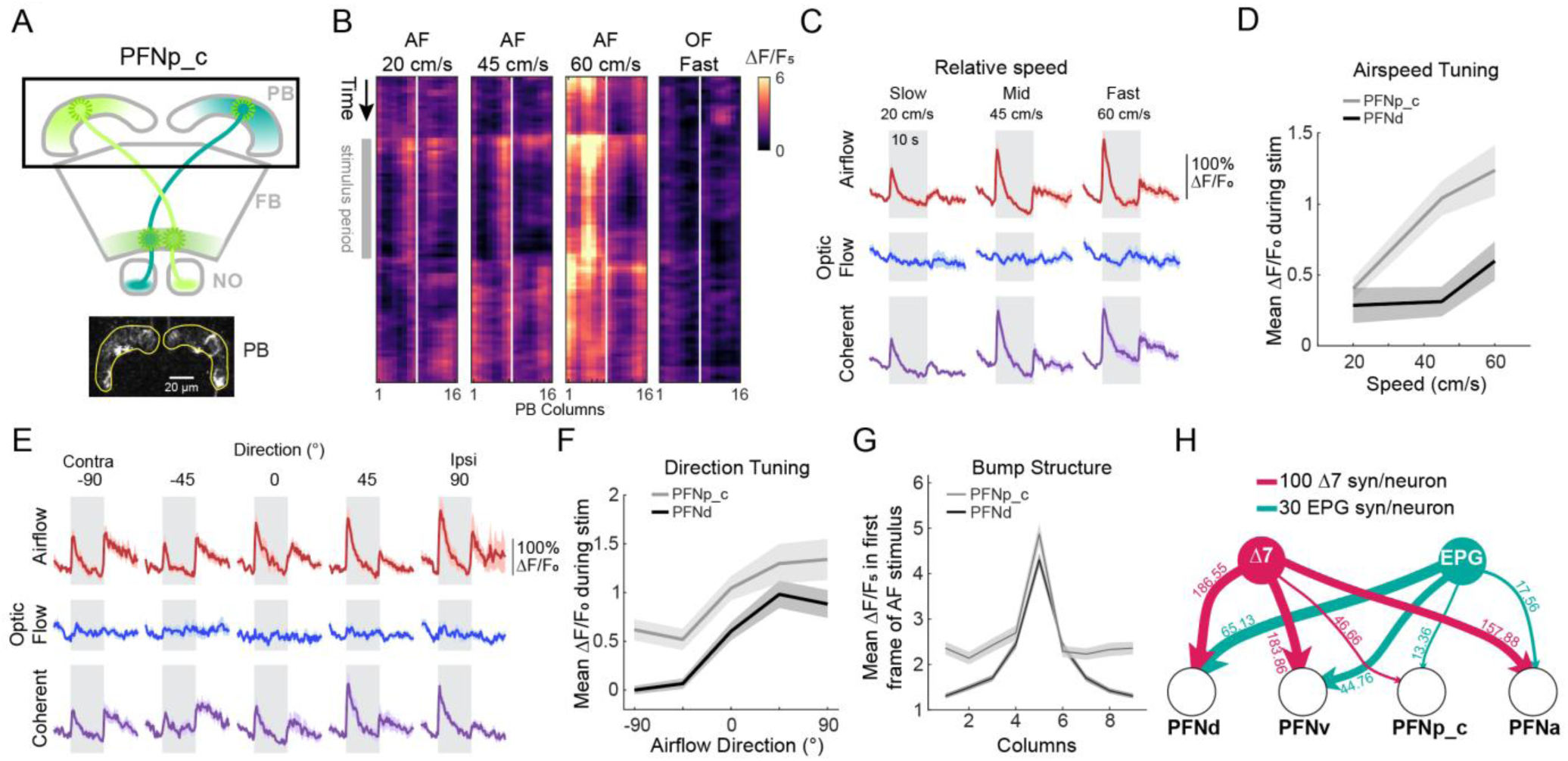
PFNp_c neurons encode airspeed and airflow direction. A. PFNp_c neuropil anatomy. Box around protocerebral bridge indicates imaging region with ROIs indicated in yellow. B. Example single-trial PFNp_c columnar calcium activity patterns in the PB to select stimuli. All data from the same fly. C. PFNp_c calcium activity in response to three stimulus speeds delivered from 0° (frontal). Traces represent mean ±SEM across flies for each PB half (N= 9 flies, 2-4 trials per fly). Gray areas indicate 10-s stimulus period. D. Airspeed tuning curves of the mean response transient (1^st^-second) amplitude for PFNp_c (light gray) versus PFNd (dark gray, same data as Supp. Fig. 1G). Lines represent mean ±SEM across flies. N=6 PFNd flies and 9 PFNp_c flies, 2-4 trials per fly, PFNp_c sdata same as in C. E. PFNp_c calcium activity in response to five stimulus directions. Coloring and shading as in C. N=7 flies, 2-4 trials per fly. F. Mean response transient (1^st^ second) amplitude of PFNp_c (light gray) versus PFNd (dark gray, same as Fig. 2E) as a function of airflow direction (contralateral is negative). Lines represent mean ±SEM across flies (N = 10 PFNd flies and 7 PFNp_c flies, n = 2-4 trials per fly). G. Bump structure in PFNp_c (light gray) versus PFNd (dark gray) as a mean fluorescent activity across columns in the first stimulus frame of every airflow trial, with fluorescent maxima aligned to column 5; column 1 is plotted twice (once on either side). Data are means across all trials ±SEM (N=10 PFNd flies and 7 PFNp_c flies, 2-4 trials per fly). H. Synaptic weights of heading-encoding neurons in the PB (EPG and Δ7) onto four PFN types. Relative line thickness and numbers indicate mean per-PFN-neuron weight calculated from hemibrain connectome (see Methods).

**Figure 3.**
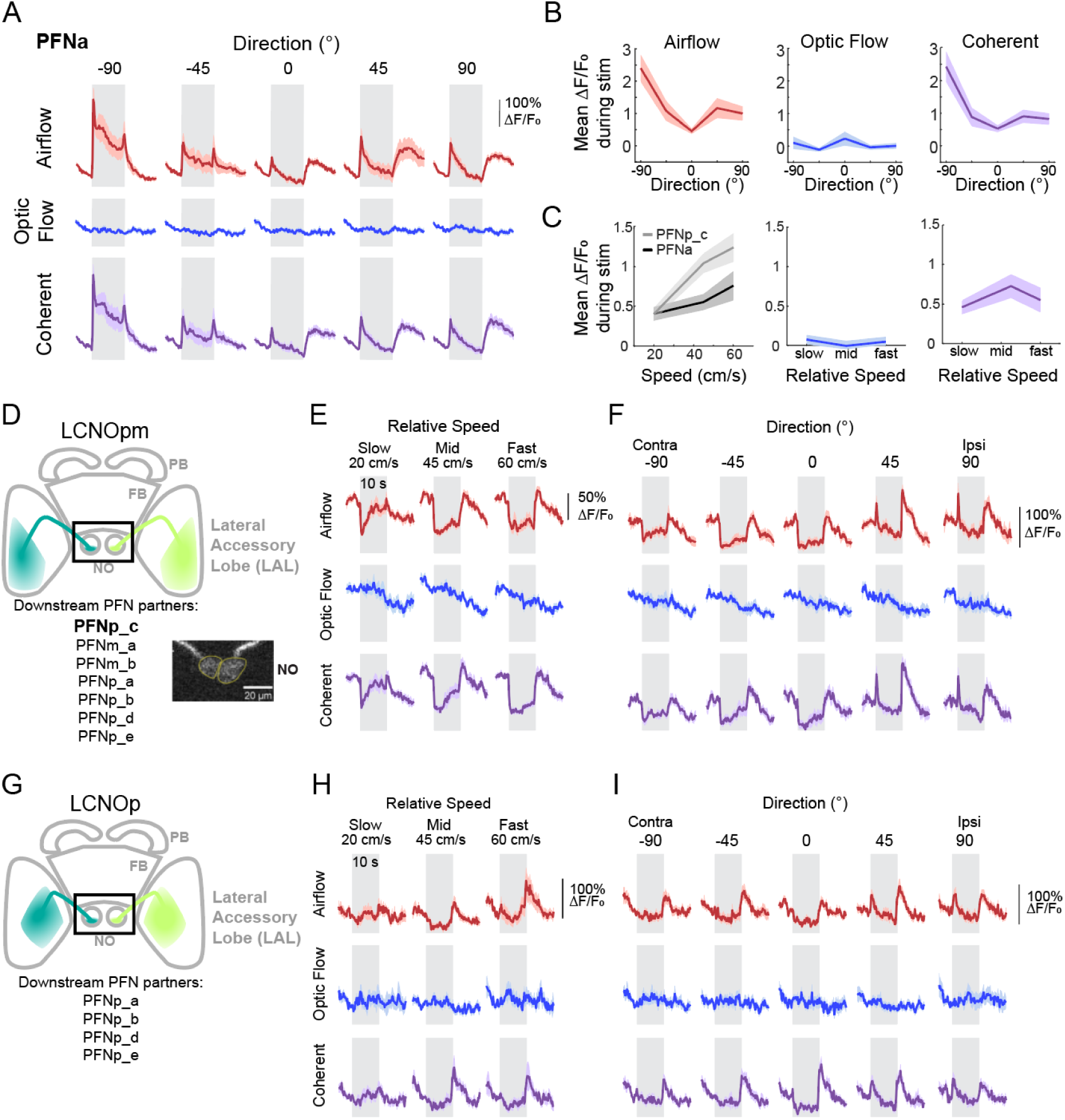
Broad encoding of airflow variables across the columnar input pathway to the FB. A. PFNa calcium activity in response to five directions of stimulus presentation; positive directions indicate stimuli ipsilateral to each PB half; negative directions indicate contralateral presentation. Traces represent mean ±SEM across flies for each PB half (N=9 flies, 3-4 trials per fly). B. Direction tuning curves for PFNa (colored traces, shading is ±SEM) for airflow (left), optic flow (middle), and the coherent stimuli (right). Data are means of the response during the entire 10-s stimulus period. C. Speed tuning curves for PFNa vs. PFNp_c (shading is ±SEM) for airflow (left), optic flow (middle), and coherent stimuli (right). PFNa data are means of the response during the entire stimulus period; PFNp_c data are means of the response transients (1^st^ second of the stimulus), same data as Fig. 2D. D. LCNOpm neuropil anatomy. Box around noduli indicates imaging region. *Inset*: single volumetric (mean z-projection) frame of imaging from noduli in LCNOpm. Left and right noduli ROIs indicated in yellow. E. Mean calcium activity in left and right noduli, combined across noduli, to three stimulus speeds delivered from 0° (N=8 flies, 2-4 trials per fly). Traces represent mean ±SEM across flies. F. Mean calcium activity in left and right LCNOpm noduli to five directions of stimulus presentation at middle speed (45 cm/s airflow, N=4 flies, 2-4 trials per fly). Traces represent mean ±SEM across flies. G. LCNOp neuropil anatomy. Box around noduli indicates imaging region. H. Mean calcium activity in left and right noduli to three stimulus speeds delivered from 0° (N=8 flies, 2-4 trials per fly). Traces represent mean ±SEM across flies. I. Mean calcium activity in left and right LCNOp noduli to five directions of stimulus presentation at middle speed (45 cm/s airflow, N=8 flies, 2-4 trials per fly). Traces represent mean ±SEM across flies.

**Figure 4.**
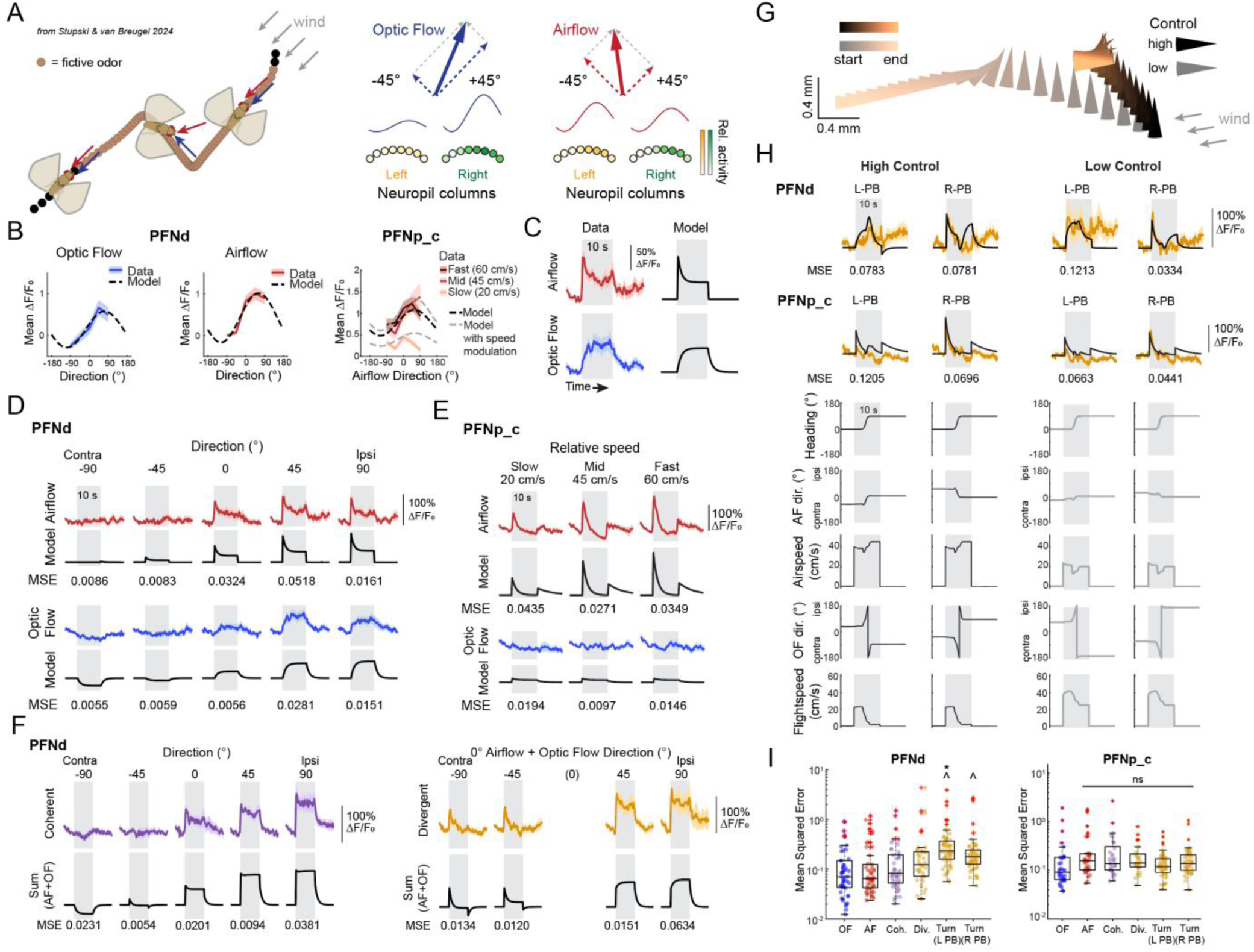
A dynamic multisensory encoding model for PFNs. A. (*left*) Example of a real active flight maneuver (*33*) that generates dynamically varying egocentric optic flow and airflow experience. (*right*) Dynamic vector coding model of PFN responses in which dynamic sensory experience is encoded by time-varying amplitudes of left and right PFN signals while bump position tracks heading as modeled previously (*24*, *28*, *31*). B. Direction tuning for optic flow or airflow was fit to a cosine function for unimodal response data for each genotype. Left and middle panels show tuning curves for PFNd (reproduced from Fig. 1G) in colored traces with overlaid fits in black dashed lines. Right panel shows direction tuning curves in PFNp_c with modulation by airspeed in grey dashed lines. C. Response dynamics for each modality were fit with an exponential decay (for airflow) or rise (for optic flow). Left panels show mean PFNd response to ipsilateral 45° airflow (*above*) or optic flow (*below*, reproduced from Fig. 1F). Right panels show modeled response dynamics (see Methods). D. Real (*top*, colored traces, reproduced from Fig. 1F) and modeled (black) responses fit to single-modality stimuli from five directions in PFNd. MSE between model and data shown below each pair of traces. E. Real (*top*, colored traces, reproduced from Fig. 2C) and modeled (black) responses fit to single -modality stimuli at three speeds in PFNp_c. MSE shown below each pair of traces. F. (*Left*) Real (*top*, colored traces, reproduced from Fig. 1F) and predicted model (black) responses to coherent multimodal stimuli in PFNd. MSE between model and data shown below each pair of traces. (*Right*) Real (*top*, colored traces, reproduced from Supp. Fig. 1A) and predicted model (black) responses to divergent multimodal stimuli in PFNd. MSE between model and data shown below each pair of traces. G. Two example trajectory simulations for delivering dynamic stimuli to tethered flies. In each, the fly performs a rightward change in heading. Each trajectory has an associated control value (high or low, see Methods). H. (*top*) Real (colored traces with shading) and overlaid predicted model (black) responses to dynamic multimodal experience of turns through wind (executed with high control (left) or low control (right)) for PFNd (above, n=10 flies, 4-5 trials per fly) and for PFNp_c (below, n=10 flies, 3-6 trials per fly). Traces are means across fly means across trials, and shading is ± SEM. MSE between model and data are shown below each trace. (*bottom*) Stimuli given to simulate experience of a right turn through a steady wind. I. (*Left*) Box and dot plots of MSE for PFNd mean response to each modality relative to model output, with each dot representing the mean MSE throughout the response and offset periods, across trials, per fly. MSEs for predicted responses to multimodal stimuli (coherent, divergent) are not significantly different (Kruskal Wallis and multiple comparisons tests, with Bonferroni correction) from MSEs for stimuli used to fit the model (OF, AF). More complex multimodal stimuli (during Turns) have larger MSEs (^ = significantly different from OF, AF, and coherent responses; * = significantly different from OF, AF, coherent, and divergent responses). (Data from n=10 flies, as in Figure 1F and 4H). (*Right*) Box plots of MSE for PFNp_c mean response to each modality (dots) per fly, relative to model output. MSEs for predicted responses to multimodal stimuli (coherent, divergent, and dynamic turn experience) are not significantly different (p = 0.1331, Kruskal Wallis and multiple comparisons test with Bonferroni correction) from MSEs for stimuli used to fit the model (OF, AF). (Data from n=7 flies, as in Figure 2E).

### Airflow speed and direction encoding in PFNp_c

We next examined sensory encoding in PFNp_c neurons (Fig. 2A), a cell type previously shown to encode airflow direction (*27*). We observed that PFNp_c neurons responded strongly and transiently to airflow, with increasing responses as a function of airspeed (Fig. 2B, C). PFNp_c responses to airflow were more transient than those of PFNd, with activity falling below baseline within 5-8 seconds of stimulus onset, and a robust offset response often following the end of the airflow stimulus (Fig. 2C). Speed tuning in PFNp_c was more robust than in PFNd (Fig. 2D), while optic flow responses were essentially absent (Fig. 2B, C, E).

In addition to airspeed tuning, PFNp_c neurons also showed airflow direction tuning (Fig. 2E). Direction tuning magnitude was comparable to PFNd (Fig. 2F), but with a positive offset, so that even the least-preferred direction (45° contralateral) produced a large transient response. Bump structure was much less obvious in PFNp_c (Fig. 2B, G) than in PFNd and we did not observe any consistent bump movement with airflow (Supp. Fig. 2A). This might reflect weaker input from the EPG compass system to PFNp_c compared to PFNd (Fig. 2H), and illustrates that not all PFNs use the same vector encoding scheme that has been described for PFNd (*24*, *31*).

### Other PFN types likely encode diverse representations of airflow speed and direction

The fly brain contains eight additional types of PFNs (*32*). Of these, PFNv was previously shown to encode optic flow (*24*) but not airflow (*27*) direction and we did not examine it further here. One other type, PFNa, can be precisely targeted using genetic driver lines and was previously shown to strongly encode airflow direction (*27*, *28*). Imaging from PFNa revealed a bimodal direction tuning curve with a small transient peak at 45° ipsilateral and a larger, more persistent response at 90° contralateral (Fig. 3A, B). Our observations are consistent with a recent study showing that hyperpolarization of PFNa by contralateral airflow drives large calcium responses mediated by T-type Ca2+ channels (*28*). We observed no response to optic flow and little tuning for airflow speed in PFNa (Fig. 3B, C). We did observe strong PFNa bump structure (Supp. Fig. 3A), as expected from its compass inputs (Fig. 2H).

The remaining six types of PFNs cannot currently be accessed with specific genetic lines (*37*). To obtain insight into what these neurons might encode, we imaged from their upstream partners in the NO, called LCNOpm and LCNOp (Fig. 3D-I). Like other LAL-NO neurons, these are predicted to be glutamatergic and inhibitory (*24*, *27*, *31*, *38*); we thus expect them to have inverted tuning relative to their downstream PFN partners, whose sensory tuning would arise through disinhibition.

LCNOpm provides input to the six PFN types in addition to PFNp_c, while LCNOp provides input to four types, a subset of those that receive input from LCNOpm (excluding PFNp_c, Fig. 3D, G) (*32*, *39*). Imaging from a specific LCNOpm line (Fig. 3D-F; Supp. Fig. 3B-D) revealed inhibition in response to airflow, similar to observations in the neuron LNOa that provides input to PFNa (*27*). The duration of inhibition grew between slow and medium airspeeds, but saturated at the highest airspeed, perhaps reflecting a floor effect from our imaging (Supp. Fig. 3D). These neurons were most strongly inhibited by airflow coming from directly in front of the fly. They were also transiently activated by ipsilateral airflow (Fig. 3F; Supp. Fig. 3C). LCNOp was weakly inhibited by airflow and showed prominent offset responses to airflow that grew with airflow speed (Fig. 3H, I). We observed no significant responses to optic flow in either LCNOpm or LCNOp (Fig. 3E, F, H, I; Supp. Fig. 3B, C, E, F). Our imaging from these two LNO neurons suggests that the additional PFN types likely also encode combinations of airspeed and airflow direction.

### Generating and validating dynamic encoding models for PFNs

We hypothesized that the multi-sensory representation of the fly’s movement through space in PFNs can be used to compute external variables such as the direction of ambient wind. To test this hypothesis, we next sought to develop a dynamic encoding model that would allow us to estimate PFN responses during real flight maneuvers (Fig. 4A, left), which are challenging to accurately replicate in head-fixed tethered flight (*40–43*).

Previous studies (*24*, *31*) have modeled the steady-state responses of PFNs as vectors, in which bump position represents a vector angle, and bump amplitude represents a vector length (Fig. 4A, right). Here we extended this framework to dynamically varying multisensory input, allowing us to simulate PFN representations to dynamic flight maneuvers. We focused our models on the three neuron types characterized here – PFNd, PFNp_c, and PFNa – and on PFNv, which has been well-characterized in other studies (*24*, *27*, *31*).

As in previous studies (*24*, *28*, *31*), we modeled PFN bump structure in half of the PB as a sinusoidal function in phase with heading (Fig. 4B). The amplitude of the bump in each hemisphere was modulated by the egocentric direction and magnitude of sensory input (optic flow and airflow, Fig. 4B). Direction tuning was specified by a cosine function peaking at ∼45° ipsilateral and fit to steady-state responses for optic flow and to transient responses for airflow (Fig. 4B). Speed tuning in PFNp_c was implemented by scaling this cosine by an exponential function of airspeed, and fitting to response transients to airflow that varied in both speed and direction (PFNp_c, Fig. 4B; Supp. Fig. 4A, B). To extend this modeling framework to dynamic inputs, we fit exponential rises (for optic flow) or decays (for airflow) to the dynamic responses we observed in our imaging (Fig. 4C). We fit the same model framework to data for PFNd, PFNp_c, and PFNa (Supp. Fig. 4A-F), with modifications to capture bimodal tuning in PFNa (Supp. Fig. 4C, E; see Methods). The resulting models closely capture the mean responses to single-modality stimulus presentation (Fig. 4D, E, I; Supp. Fig. 4A-C).

To validate our model we took two approaches. First, we asked whether models fit to single-modality data could be used to accurately predict responses to multimodal input. Based on our observations in PFNd, we modeled responses to multimodal stimuli as a sum of the responses to optic flow and airflow alone (see Methods). We predicted responses for the coherent and divergent stimuli we delivered in our initial experiments (Fig. 4F), as well as for a set of “slip” experiments (Supp. Fig. 4G,H) in which airflow and optic flow were dynamically and coherently shifted by 45° midway through stimulus presentation. Overall, we found that our model provided a good fit to these multi-sensory responses, as quantified by mean squared error (MSE) (Fig. 4I; Supp. Fig. 4H). Real responses were more transient than predicted responses suggesting a greater weighting on airflow input, as well as an additional slow decay, especially during the longer slip stimuli (Supp Fig. 4I). As a further test of our model, we performed a new set of experiments, in which we simulated the experience of a flying fly turning in the presence of wind. We used proportional valves to generate dynamic airflow speeds over time, and rotated the airflow stimulus to simulate a change in airflow direction (Fig. 4G-I, Methods).

Due to physical hardware constraints, these rotations were somewhat slower than what a fly making a true saccadic flight turn would experience. Our model also provided a good fit to these more complex and naturalistic stimuli, although optic flow responses in this dataset were somewhat weaker than predicted (Fig. 4H, Supp Fig. 5). We conclude that our dynamic encoding model allows for reasonable simulation of PFN responses during complex flight maneuvers (Fig. 4I).

**Figure 5.**
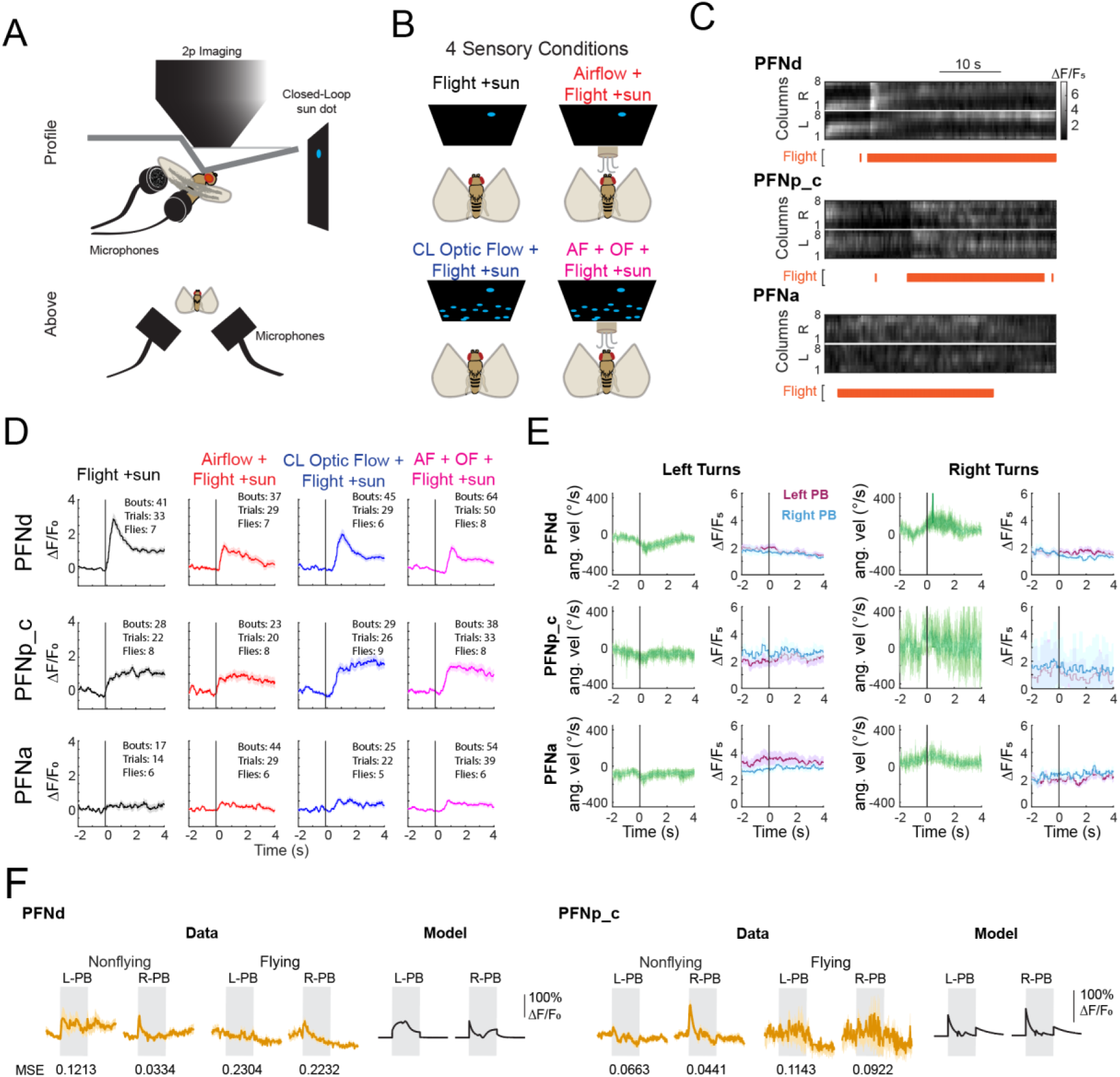
PFNs remain active during flight, do not encode flight maneuvers in tethered flies, and mostly respond as predicted to dynamic stimuli during flight. A. Paradigm for recording flight behavior during 2-photon imaging. Microphones positioned behind each wing measure left and right wingbeat amplitude and frequency. B. Four closed-loop sensory conditions activated during flight: *top left*, no airflow or optic flow, only sun dot in closed loop; *top right*, sun dot in closed loop and airflow from 0° (front of the fly) only when fly is flying; *bottom left*, sun dot in closed loop and forward optic flow experience in closed loop with turning behavior and flight; *bottom right*, sun dot in closed loop with airflow from 0° and forward optic flow in closed loop with behavior. C. Representative calcium activity in PB columns of PFNd (*upper*), PFNp_c (*middle*), and PFNa (*lower*). For each calcium data plot, the recorded flight state (see Methods) for the trial is plotted below. D. Mean calcium activity in the PB in each of the three PFN types across the four conditions in A. Flight bout begins at time = 0 s (black vertical line). All flight bouts longer than 4 s were included. Traces are means across bouts ±SEM (n=37-64 bouts across 29-50 trials in 7-9 PFNd flies, n=23-38 bouts across 20-33 trials in 8-9 PFNp_c flies, n=17-54 bouts across 14-39 trials in 5-6 PFNa flies). E. Mean calcium activity around left or right turns for each PFN type (see Methods). n=31 left and 16 right turns across 16 trials in 7 flies for PFNd; n=14 left and 3 right turns across 8 trials in 5 flies for PFNp_c; n=22 left and 9 right turns across 13 trials in 6 flies for PFNa. Data drawn from same set as C. F. Mean calcium activity for the left and right halves of the PB during dynamic sensory experience of non-saccadic turn through ambient wind, for PFNd (left) and PFNp_c (right). For each set, the leftmost two traces (colored trace is mean; shading = ±SEM) for each are from nonflying flies, same data as Fig. 4H; the middle two traces are the responses to the same stimuli in flying flies. The modeled responses are the black traces on the right. For the data from flying flies, PFNd had 12 bouts of constant flight for the duration of the stimulus, drawn from 5 flies; PFNp_c had 6 bouts of constant flight for the duration of the stimulus, drawn from 3 flies. MSE reported below each data trace compares to the model trace.

We considered whether, in addition to airflow and optic flow, PFNs might encode self-motion through efference copy or proprioception during flight. Indeed, imaging results from walking flies suggest that they may receive proprioceptive input about self-motion (*31*). To examine encoding of flight maneuvers we imaged from tethered flying flies, and measured wing flapping behavior using two microphones positioned behind each wing with four sensory feedback conditions (Fig. 5A,B). We observed that PFNd and PFNp_c showed increased activity during flight behavior, while PFNa activity remained robust but did not increase (Fig. 5C). Experience of frontal airflow minimally inhibited PFNd and PFNp_c activity during flight (Fig. 5D).

We then asked whether flight turn information (perhaps through efference copy) might also be represented in PFNs. To address this question, we detected fictive flight turns as differences between the two microphone signals that crossed a threshold (see Methods), and plotted PFN activity associated with either left or right turns (Fig. 5E). We observed no significant signals related to fictive turns, arguing that efference copy of these maneuvers is not encoded by these PFNs, at least in tethered flight when halteres are not activated. We also tested the predictions of our model by measuring responses to simulated flight turns in flying flies (Fig. 5F, Supp. Fig. 5B-C). Though we saw some reduction in PFNp_c ΔF/F response amplitudes, likely due to higher baseline activity from flight-induced activation, the dynamics of the responses in both types broadly matched the model predictions (Fig. 5F, Supp. Fig 5B,C). We conclude that PFNs are active during flight and that they encode primarily optic flow and airflow, as characterized here, and heading, as characterized previously (*24*, *31*), although it is possible that input from the halteres, which are activated during free flight turns, could be encoded by these neurons (*44*).

### Ambient wind direction is observable from PFN activity during saccadic flight turns

Armed with our dynamic encoding model, we set out to ask whether the representation of self-motion in PFNs is sufficient to decode ambient wind direction in flight, as has previously been shown for raw sensor measurements (*10*, *11*). We first simulated responses of the PFNs to the reconstructed sensory experience of real flight trajectories recorded in a wind tunnel (*33*). In these recordings, freely flying flies expressing CsChrimson in *orco+* olfactory receptor neurons were subjected to a flash of red light, providing a fictive odor stimulus. As reported previously, fictive odor stimulation often led to two turns: a random-direction “anemometric” turn and deceleration in the first 50-100 ms that we hypothesize allows the fly to estimate wind direction, followed by an upwind “anemotactic” turn ∼100-150 ms later (Fig. 6A). Using our dynamic encoding model, we simulated the responses of four PFN types to real flight trajectories, providing a rich but concise neural representation of self-motion in wind (Fig. 6B, C).

**Figure 6.**
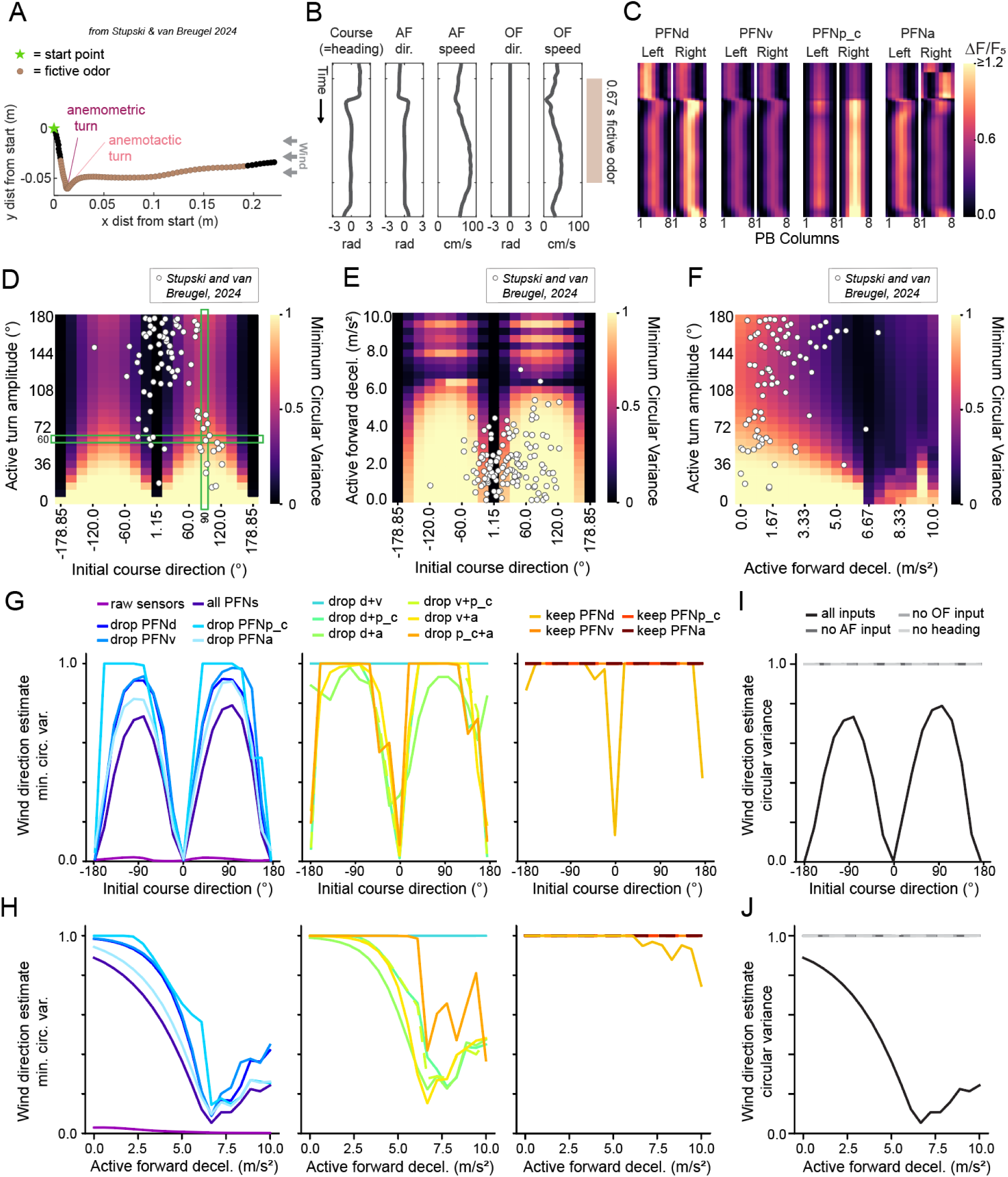
Wind direction during flight is observable from PFN representations. A. Real horizontal trajectory of a freely flying fly in a laminar wind tunnel in response to fictive optogenetic odor (gold). Data from (*33*). The fly is initially moving crosswind, then decelerates and makes a turn away from upwind in response to the fictive odor (anemometric turn, burgundy), and finally turns upwind shortly after (anemotactic turn, pink). Sample rate is 100 Hz. B. Measured course direction (assumed heading-aligned) and estimated egocentric airflow, and egocentric optic flow experience during the trajectory in A . C. Predicted activity in four PFN types using the models fit in Fig. 6 and the inputs shown in B. D. Heatmap of wind direction estimate error circular variance as a function of initial course direction and turn amplitude. Error computed from observability analysis of simulated 100-ms saccadic flight trajectories with 2.5 m/s^2^ forward deceleration (see Methods). White data points show course directions and turn magnitudes of real flight trajectories from (*33*). Green outlined regions indicate parameters held constant for G-J. E. Same as D but for simulated saccadic flight trajectories of turn amplitude 20°, varying across forward deceleration magnitudes. F. Heatmap of wind direction estimate error circular variance as a function of forward deceleration and turn amplitude, all with initial course direction of 90° and wind from 0°. G. Effects of sensor dropout on wind direction estimate error circular variance for 60°-turn saccades with 2.5 m/s^2^ deceleration across various initial course directions. Left: results using raw sensors, all PFNs (as in D,E,F), or removing single PFN types. Middle: effects of removing pairs of PFN types. Right: effects of removing any three PFNs, leaving one as the sole sensor set. H. Same as J, but reporting observability from 60°-turn trajectories with initial course of 90°, varying across forward deceleration magnitude. I. Effects of dropping heading, AF, or OF information on wind direction estimate error circular variance, varying across initial course directions.. J. Same as L, but reporting observability from 60°-turn trajectories with initial course of 90°, varying across forward deceleration magnitude.

We next applied a nonlinear empirical observability analysis (*10*) to ask whether ambient wind direction can be decoded in principle from these PFN representations. Derived from Fisher information (*45*), observability can be used to determine what unknown information can be estimated from a set of sensory measurements (*8*, *46–48*), including those that vary over time (*10*) (see Methods). Our observability analysis pipeline consisted of three steps. First, we simulated sensory variables available to a fly during 100-ms saccade-like turn trajectories with both deceleration and a sharp change in course direction (Supp. Fig. 6A-C) using model predictive control (*33*, *49–51*). These trajectories were defined by flight speed, course direction, angular velocity, ambient wind direction, ambient wind speed, and altitude. For simplicity, in these simulations we assumed the fly was facing its direction of motion such that heading and course direction were equivalent. Because the heading angle simply defines the coordinate frame, it has essentially no impact on observability (*52*), thus we do not expect this simplification to impact our results. From these, we produced time courses of the fly’s (global-coordinate) heading, egocentric airflow direction and speed, and egocentric optic flow direction and speed, and fed these five sensory experience variables into our PFN activity models. We note that optic flow speed is not encoded by PFNs in our data and so effectively disappears at this stage. We then use observability to compute a metric for the minimum possible unbiased reconstruction error circular variance for the wind direction, given a set of PFN neural responses as measurements (see Methods for specific details). A small error variance indicates that wind direction can be accurately decoded (more observable), whereas a large error variance means it cannot (less observable).

Our analysis revealed that wind direction observability depends strongly on parameters of flight trajectories including initial course, turn amplitude, and deceleration (Fig. 6D-F). We found that wind direction was observable (reconstruction error variance < 0.5) for most saccadic turns except for small turns (<∼45°) with decelerations that maintained forward motion (<∼6 m/s^2^) (Fig. 6D-F). Most real measured active flight maneuvers— both turns and decelerations— (*10*) (white dots in Fig. 6D-F) fell within the observable range, supporting the idea that active flight maneuvers, combined with biological encoding of sensory feedback, are sufficient to decode the ambient wind direction under natural conditions. By eliminating individual PFN types or groups from our analysis, we asked how different PFN types contribute to the observability of wind direction (Fig. 6G,H). We found that removing a single PFN type produces little change in the estimate error variance (Fig. 6G and H, left), likely because most information is redundantly represented across multiple PFN types. The exception is PFNp_c, which is the only PFN type that encodes airspeed in our model. Removing both PFNd and PFNv, which are the only two PFNs that encode optic flow, produced a large increase in reconstruction error. Eliminating all optic flow, or all airflow information, or removing heading encoding (similar to what we expect if EPG neurons are silenced) all drastically increased the error in wind direction estimation (variance near or at 1) (Fig. 6I,J). We conclude that encoding of heading, airflow direction, airflow speed, optic flow direction all contribute to the ability to decode wind direction, but that silencing individual PFN types may not produce large effects because of redundant encoding of movement parameters across types.

### Ambient wind direction can be decoded from PFN representations by a biologically-inspired artificial neural network

Observability analysis allows us to determine whether wind direction can be decoded from PFN representations, but not how. To obtain insight into how wind direction might be decoded by biological circuits, we constructed and trained a set of simple feedforward artificial neural networks, based on motifs from the fly connectome (Fig. 7A). In our networks, units representing PFN activity provide input to one or more hidden layers. In order to capture changes in the PFN representation that arise from wind or active movement, the input layer represented PFN activity at two time-points: 0 and 12 ms in the past. The hidden layer(s) then provide input to an 8-unit output layer modeled on local neurons of the CX. We reasoned that an estimate of wind direction in flight would be represented as a sinusoidal bump of activity across this output array with the position of the bump representing allocentric wind direction (*28*). We constructed and trained three ANNs with the same input and output structure, but which varied in number of hidden layers (1 or 2), hidden units per layer (36 or 18), and training epochs (500 or 1000; see Methods, Supp. Fig. 7A).

**Figure 7.**
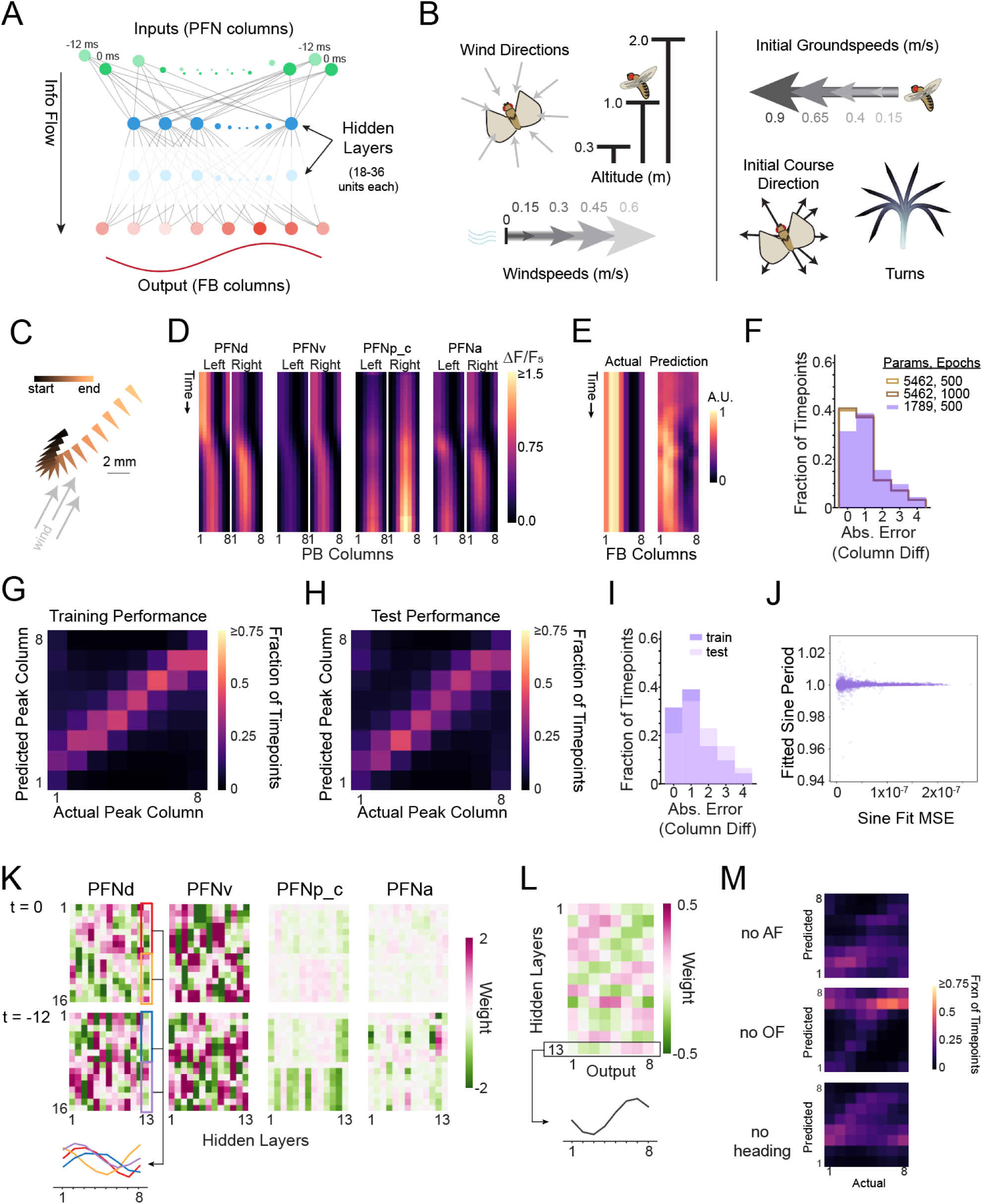
PFN encoding is sufficient for a compact feedforward ANN to estimate wind direction. A. General structure of the trained and tested ANNs. Input units correspond to calcium intensity in individual PB columns for PFNd, PFNv, PFNp_c, and PFNa at two time points. Output units correspond to column calcium intensity values in a putative allocentric wind representation in a local neuron type within the FB. B. Parameters for the flight trajectories underlying sensory experience in the training dataset. C. An example trajectory from the training dataset. D. Predicted columnar PFN activity during the trajectory shown in C. E. Actual and predicted columnar representations from the ANN with the fewest trainable parameters during the trajectory shown in C. F. Distribution of circular absolute difference in peak column position for three ANNs varying in number of hidden layers and of training epochs. The most compact ANN (shaded purple) has only one hidden layer and the fewest trainable parameters. No significant difference between any pair of distributions, by mean of resampled Kuiper tests (p-values>0.42; see Methods). G. Performance of the most compact ANN on the training dataset as a 2D histogram showing the distribution of output columns housing the peak of the output sinusoid across all the timepoints in the trajectories of the training dataset. The distributions have been normalized to the number of actual peak column positions (x-axis) so that each column of the heatmap sums to 1.0. H. Performance of the most compact ANN on the pseudorandom test dataset as a 2D histogram of predicted vs. actual output peak columns, as in G. I. Distribution of absolute circular column difference between predicted and actual wind direction sinusoid peak column positions for the training and test datasets using the most compact ANN. No significant difference between the distributions, by mean of resampled Kuiper tests (p>0.917). J. Estimated period parameter values for sinusoid fits to every output set from the ANN, plotted against the mean squared error of the sinusoid to the ANN’s output set. K. Learned weights from inputs to hidden layer of the most compact ANN, separated by input type (PFN type and timepoint). Below, the values of the PFNd input weights onto the 13^th^ hidden layer unit are plotted to demonstrate sinusoidal shape. Each set of 8 units pertaining to a timepoint and the columns of half of the PB generate sinusoidal weight structure. L. Learned weights from hidden layer units to outputs of the most compact ANN. Below, the values of the 13^th^ hidden layer unit weights onto the 8 output units are plotted to demonstrate their sinusoidal shape. M. 2D histograms showing the performance of the most compact ANN on the pseudorandom test dataset after removing PFN encoding of key sensory information (airflow, optic flow, or heading).

We next constructed training and test datasets designed to mimic PFN representations of sensory feedback in the presence of a fixed ambient wind direction under a wide range of flight conditions. For each 100-ms trajectory, we selected one of 8 ambient wind directions, represented as sinusoids in the output layer with different phases. For each wind direction, we then simulated trajectories with diverse altitudes, wind speeds, initial course directions and speeds, for a total of 30,720 trajectories (Fig. 7B). Each trajectory contained one turn of varying magnitude, accompanied by a deceleration. The full trajectory (*e.g.,* Fig. 7C) was calculated using model predictive control (see Methods). For these simulations, we included an estimate of heading that naturalistically deviated from course direction predicted by a network trained on flights performed by real flies with accurately measured body orientations and course directions (*3*, *53*). The resulting PFN representations of sensory feedback were then computed using our PFN encoding model (Fig. 7D). These trajectories were used to train the ANN to map pairs of PFN representations to a single sinusoidal output pattern representing the ambient wind direction for each time point in the trajectory (Fig. 7D,E). A separate test set of 5000 trajectories was generated using parameters randomly selected from reasonable ranges for each variable (see Methods).

Of the three trained ANNs (Supp. Fig. 7A), we found that the one with the fewest parameters (a single hidden layer with 18 units) performed similarly to the larger ones on the training dataset (Fig. 7F,G). The performance of this model on held out test data was very similar to its performance on the training data (Fig. 7H,I; Supp. Fig. 7B). The learned weights onto this hidden layer, and from the hidden layer to the output layer, had a sinusoidal organization (Fig. 7J-L), similar to patterns of weights observed from PFN types onto local neuron types (hΔ and vΔ), and from local neuron types onto other local neuron types in the CX (*32*). We further compressed this model by eliminating hidden units with very small input weights (Fig. 7K, Supp. Fig. 7C,D, see Methods) to yield a model with 13 hidden units. This had no effect on the sinusoidal weight patterns.

To probe the contributions of different modalities to ANN performance, we remade the test dataset absent PFN responses to either airflow, optic flow, or heading and again used the network to get predictions of wind direction (Fig. 7M). The lack of any of these variables greatly impaired the network’s performance. We conclude that a simple feedforward ANN, with connectivity motifs similar to those observed in the fly CX, is capable of decoding ambient wind direction from realistic PFN representations of heading and multi-modal sensory feedback.

### The windspeed-to-groundspeed ratio defines two solution regimes in both the ANN and real flies

The strong performance of the ANN led us to wonder what algorithm it uses to decode ambient wind direction. Examining the training data for each allocentric wind direction (corresponding to one output configuration), we noticed two solution regimes that depended on the ratio of windspeed to groundspeed (Fig. 8A). At high windspeed-to-groundspeed ratios, there is a simple linear relationship between course direction and airflow direction (Fig. 8A: red points). This is because, in this regime, the ambient wind vector and the airflow vector are similar in both direction and magnitude (Fig. 8B). In contrast, when the windspeed-to-groundspeed ratio is less than 1 (Fig. 8A: green and blue points), mechanosensory feedback more strongly reflects self-generated motion rather than wind (Fig. 8B).

**Figure 8.**
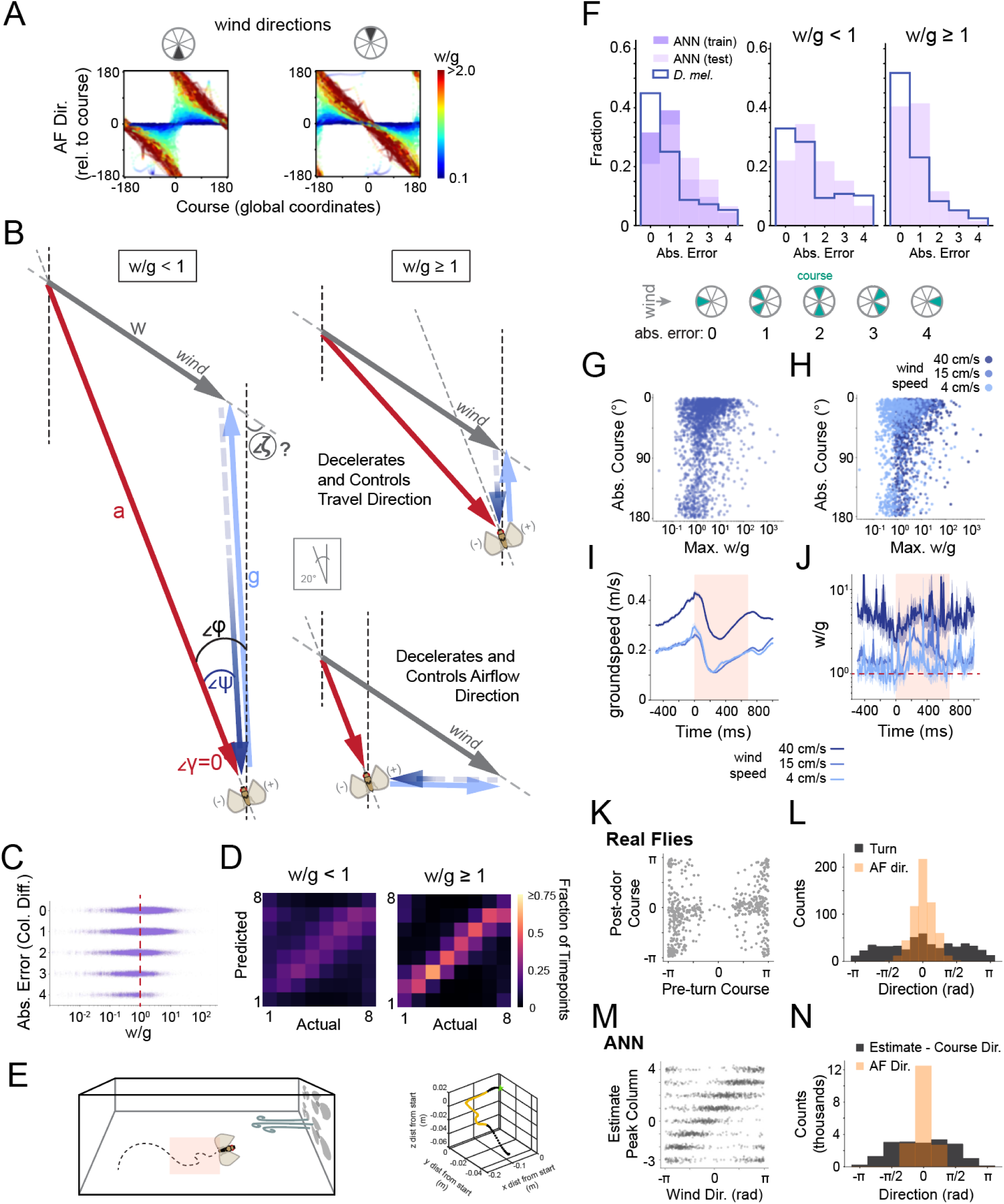
The windspeed-to-groundspeed ratio segregates trivial from high-difficulty wind direction solutions for real flies and the PFN-like ANN. A. Scatter plots of the relationship between course direction (x axis) and egocentric airflow direction (y axis) across the randomized test dataset for the ANN. Each plot contains data from a 45° range of wind directions as indicated in the pie chart. Each dot is a timepoint within a trajectory; the color of each dot indicates the windspeed-to-groundspeed ratio at that timepoint. B. When the ratio of windspeed to groundspeed is less than 1 (*left*) the direction of airflow experience is largely driven by feedback from flight. When this ratio is greater than one (*right*), a change in direction or speed leads to new wind triangles that share a common leg (ambient wind vector). Controlling optic flow (*above*) or airflow (*below*) direction causes the other modality vector to become more aligned with wind, allowing for wind direction estimation. C. Scatter plot of the difference in peak column position between the actual wind direction representation and the ANN’s predicted output for every input, plotted against the windspeed-to-groundspeed ratio for that input. D. 2D histograms showing the performance of the most compact ANN on the pseudorandom test dataset, for low (*left*) and high (*right*) windspeed-to-groundspeed ratios. E. Schematic of real flies navigating a wind tunnel (*left*) with fictive odor triggered by flight through a cubic space (pink shading), and an example trajectory (*right*) from (*33*). F. (*Left*) Comparison between the distribution of wind direction estimation error for the ANN (same data as in Fig. 7J) and for real flies. (*Middle* and *right*) Same comparison separated into low (left) and high (right) windspeed to groundspeed ratio. Course direction differences from upwind (as ‘absolute error’) were binned as shown in the key (*lower*). All comparisons tested with mean of resampled Kuiper tests; p-value ∼0.001 for real flies vs. ANN training performance, real flies vs. ANN test performance, real flies vs. ANN test performance in w/g<1 regime, and real flies vs. ANN test performance in w/g≥1 regime. G. Relationship of maximum windspeed-to-groundspeed ratio achieved during anemometric flight maneuvers (deceleration) in real flies (x axis) versus absolute value of course direction after anemotactic turn (y axis; 0° represents upwind). Key in F defines how absolute value of course was binned to compute error in bottom plot. H. Relationship of maximum w/g during anemometric flight maneuvers, same data as G with dots colored by ambient windspeed. I. Groundspeed (m/s) over time during odor experience in real flies in three different wind conditions. Same dataset as G and H. Pink shading indicates duration of fictive odor experience. J. Windspeed-to-groundspeed ratio (w/g) over time in three different wind conditions. Same dataset as G-I. Pink shading indicates duration of fictive odor experience. K. Scatter plot of the course for real flies before (x axis) and after (y axis) an anemotactic turn, for all trajectories where course direction before the turn differs from upwind by more than π/4 radians and windspeed-to-groundspeed < 1. One point represents one contiguous recorded trajectory. L. Comparison of the distributions of the anemotactic turn angle (charcoal) and the direction of airflow relative to course direction experienced before the anemotactic turn (pale orange) for same trajectories as in K. Statistically significant (p-value ∼ 0.001) by mean of resampled Kuiper tests. M. Scatter plot of the true wind direction (x axis) and the column location of the ANN’s estimate of the wind direction for inputs at the seventh timepoint (pre-turn) in each trajectory in the test dataset, with course direction different from upwind by more than π/4 radians and windspeed-to-groundspeed ratio < 1. N. Comparison of the distributions of the possible turn (as wind direction estimate minus current course direction, in charcoal) for the same inputs as M, and their associated airflow direction relative to course (pale orange). Statistically significant (p-value ∼ 0.001) by mean of resampled Kuiper tests.

These observations led us to ask whether the behavior of the ANN depends on this ratio. We plotted the performance of the ANN for trajectories with windspeed-to-groundspeed ratios greater than or equal to one, or less than one (Fig. 8C,D, Supp. Fig. 9A-B). We found that performance was much better for the high windspeed-to-groundspeed ratio regime (Fig. 8C,D), although performance in the low windspeed-to-groundspeed regime was still better than chance (p<0.0001, Rayleigh’s).

We next wondered how the performance of the ANN compares to the performance of real flies, and whether real flies also exhibit a greater ability to determine ambient wind direction when windspeed is higher than groundspeed. To address this question, we collected a new dataset of flight trajectories in response to optogenetic activation of olfactory receptor neurons (Fig. 8E). In previously published data (*33*), flies expressing CsChrimson in the olfactory receptor neurons (ORNs) received a brief light pulse to simulate appetitive odor experience in the wind tunnel. Using the same paradigm, we collected new data at lower wind speeds and combined this with the previously published set for three total wind speeds: 40cm/s, 15 cm/s, and 4 cm/s. We measured the flies’ course direction 135-400 ms after optogenetic activation of ORNs and divided trajectories into two groups depending on whether the maximum windspeed-to-groundspeed ratio achieved during the fictive odor was greater than or equal to one. Similar to our observations with the ANN, we found that real flies performed significantly better when the windspeed-to-groundspeed ratio was greater than or equal to one (Fig. 8F-H), although performance was significantly better than chance (p<2.87e-77, Rayleigh’s) even at low windspeed to-groundspeed ratios. Real flies performed better than the ANN across both regimes (p-values ∼ 0.001, Kuiper’s test with resampling; Fig. 8F). These effects can also be seen by plotting the fly’s final course direction, or its error relative to upwind, as a function of maximum windspeed-to-groundspeed ratio (Fig. 8C,G,H, Supp. Fig. 9C).

In a previous study (*33*), we found that flies often decelerate in response to optogenetic activation of olfactory receptor neurons, which should increase the windspeed-to-groundspeed ratio while windspeed is stable. We therefore analyzed changes in groundspeed evoked by optogenetic activation in this new dataset. We found that optogenetic activation led to a brief and stereotyped deceleration that was similar across wind speeds (Fig. 8I, Supp. Fig. 9C-F). As a result of this deceleration, the windspeed-to-groundspeed ratio increased following fictive olfactory stimulation, placing the majority of trajectories in the regime where windspeed-to-groundspeed is greater than or equal to one (Fig. 8J).

Finally, we asked to what extent flies and the ANN used egocentric airflow direction as a proxy for ambient wind direction in the low w/g regime, and whether they are likely using similar or different algorithms to detect deviations from this estimate. We first gathered trajectories from real flies whose windspeed-to-groundspeed ratio was less than 1 and whose course direction before the anemotactic turn was at least π/4 radians away from upwind, to ensure the flies made an active decision to turn.

We show that the majority of these trajectories result in upwind course following the anemotactic turn (Fig. 8K). We then show that, for many trajectories, flies make turns much larger than the magnitude of egocentric airflow direction by plotting these distributions and running resampled Kuiper tests on the resulting data (Fig. 8L, p-value ∼ 0.001; see Methods). This indicates that they do not solely use airflow direction to estimate wind direction.

To perform the same test on the ANN, we first plotted the columnar phase of the ANN’s output against the actual wind direction, for early-trajectory inputs where course direction was not upwind and w/g < 1. Fig. 8M demonstrates that for this data subset many of the estimates have low error (densities along the diagonal). To compare ANN function to real fly behavior, we computed a potential “turn” that could be generated from the ANN’s estimate: the difference between its wind direction estimate and its current course direction. We reasoned that if the ANN were to use airflow direction as a proxy for wind direction at this early timepoint in the trajectory, then the distribution of egocentric airflow directions and the distribution of these turn angles would match. We directly compared these using Kuiper’s test with resampling, and found the distributions are significantly different (Fig. 8N, p-value ∼ 0.001). We conclude that when w/g < 1, neither real flies nor our ANN consistently estimate wind direction using only airflow direction. Both systems must therefore perform a computation with multiple sensory variables encoded by PFNs.

## Discussion

### A multi-sensory representation of self-motion in the columnar input pathway to the insect navigation center

To navigate through space, the brains of both vertebrates and invertebrates construct internal representations that allow them to generalize across sensory inputs and make spatial inferences (*20*, *54*, *55*). For example, the compass system of flies integrates both visual (*56–58*) and mechanosensory (*36*) cues about heading to produce an “abstract” representation of heading direction (*59*). This representation allows the fly to associate a particular direction with punishment (*60*) or a wind direction with visual landmarks (*61*). Similarly, place cells of the rodent hippocampus generate an abstract representation of space thought to allow the animal to navigate towards remembered locations based on distal cues (*62–64*). Abstract representations also emerge in artificial neural networks trained to perform spatial tasks (*65*, *66*). What kinds of abstract internal representations each nervous system builds, and what computational function these representations serve are currently topics of intense research interest.

In vertebrates, several regions of the brain have been shown to integrate multi-sensory cues that signal self-motion. For example, neurons in cortical areas MSTd, VIP, and 7a integrate both optic flow and vestibular cues that signal translational self-motion (*22*, *67*, *68*). Single neurons can be tuned either to the same direction (congruent cells) or to opposing directions (opposite cells) (*22*). Population activity provides a good fit to the animal’s reported direction of self-motion (*22*). Multi-sensory input from these areas, especially 7a, are thought to provide indirect input to spatially tuned neurons in the retrosplenial cortex and medial entorhinal cortex (*69*, *70*), which support allocentric computations of location in space (*71*), in part by integrating self-motion information (*22*, *72*), but the precise pathways involved in this transformation are not known.

Here we show that the LNO-PFN pathway that provides input to the insect navigation center builds a multi-sensory representation of self-motion. In insects, optic flow is computed by T4/T5 neurons of the optic lobe (*73*, *74*) and integrated by lobula plate tangential neurons (*75*) to produce representations of typical flight maneuvers (*12*). Mechano-sensory neurons in the antenna, known as JONs, respond to antennal movement produced by both self-motion and ambient wind (*76–79*), and may function as a kind of vestibular system in some insect species (*30*). Airflow direction can be decoded from differential displacements of the two antennae (*36*, *76*), and is likely carried to PFNs by a pathway involving the AMMC, WED, and LAL (*27*, *36*). Strikingly, multi-sensory PFNs (PFNd) are tuned to optic flow and airflow signals from the same direction, 45° ipsilateral to the protocerebral bridge innervation of each neuron. Moreover, we find that the visual and mechano-sensory responses of these neurons exhibit distinct time courses, consistent with the known sensitivity of each sensory system. Visual responses are slow and sustained, reflecting the slow but accurate nature of visual self-motion direction signals, while mechano-sensory responses are fast and more transient, reflecting the more transient nature of mechano-sensory cues about self-motion. We found that PFNd combines multi-sensory direction information approximately linearly, regardless of the degree of coherence of the two stimuli. This suggests a role for PFNd activity in reflecting self-motion direction, even in the absence of one of the modalities, e.g. if the antennae were damaged or if the fly were flying in darkness.

To determine its location, an animal must keep track of movement speed as well as direction. Previous studies of speed coding in PFNs have yielded conflicting results. PFNd and v neurons did not show tuning for optic flow speed (*24*) but did show velocity tuning for walking (*31*). Here we measured optic flow and airflow speed tuning across multiple PFN and LNO types. We did not observe any tuning for optic flow speed, but did observe robust tuning for airflow speed in both PFNp_c and two LNO types: LCNOpm and LCNOp. Together these data suggest that mechano-sensory cues (airflow and proprioception), rather than optic flow, might be used to estimate speed in the fly navigation center. Curiously, all of the speed-tuned responses we observed were transient, responding either to the onset or offset of airflow, or both. This suggests that these responses may encode airflow (or self-motion) acceleration. Acceleration-tuned neurons have been observed at many layers of the vertebrate vestibular system (*80*).

A limitation of our study is that there are currently six PFN types that cannot presently be individually targeted with genetic reagents. Imaging from the upstream partners of these neurons, called LNOs, revealed airflow speed and direction tuning, but no optic flow responses. It is therefore likely that some of these “dark PFNs” may encode other functions of airflow (or mechano-sensory self-motion feedback). For example, they might encode these variables over different dynamic ranges or integrate them over different timescales. In addition, many of these PFNs may encode proprioceptive cues about self-motion. Although we did not observe encoding of turning maneuvers in tethered flight, natural flight strongly activates the halteres (*81*), which are known to provide input to the navigation center in other fly species (*44*). Activity related to walking has also been recorded in some PFNs (*31*). Future studies examining the integration of airflow and optic flow with proprioceptive information from the legs and halteres will allow us to understand how PFNs as a population encode self-motion.

### PFN representations are sufficient to compute ambient wind direction when combined with active sensing

What is the function of the multi-sensory representation in PFNs? One established function of optic flow-tuned neurons is to enable a transformation from egocentric to allocentric coordinates (*24*, *31*). It has been shown theoretically that PFNa could in principle compute allocentric wind direction through a similar vector summation algorithm in walking flies (*28*). However, these computations do not explicitly require multi-sensory integration. Here we propose a third possible function for the PFN representation: allowing the computation of ambient wind direction while in flight. Wind direction is an essential cue for flying flies (*5*, *33*), similar to flow direction for fish (*82*, *83*). Catching the wind can power dispersal over longer distances, while upwind and crosswind orientation facilitate different stages of olfactory navigation (*84*). In flight, self-generated movement and ambient wind sum together producing ambiguous visual and mechanosensory sensory feedback. However, optogenetic free flight experiments argue that flies can estimate the wind direction in flight by making active “anemometric” turns or decelerations (*33*). Upwind turns in this paradigm occur on very short timescales (∼50 ms) (*33*) that are too slow to be mediated by visual feedback (*34*, *35*), arguing that they rely instead on an internal estimate of the ambient wind direction. Mathematically, a flying fly can compute the ambient wind direction if it has access to sufficient angles or lengths of the wind triangle (*8*, *11*). Here we showed that PFNs encode one angle and one length of this triangle: airflow and optic flow direction relative to the fly, and airflow speed. Notably, we observed little encoding of optic flow speed for our stimuli. This might reflect the fact that groundspeed is challenging to estimate accurately from optic flow speed (*85*, *86*). A previous study in flies also observed little encoding of optic flow speed using starfield stimuli similar to ours (*24*), while a study in bees found optic flow speed encoding using grating stimuli in TN neurons (*26*), homologous to LNO neurons in fly. These differences might reflect differences between species, stimuli, or neurons targeted.

To test the sufficiency of PFN representations for decoding wind direction, we generated dynamic encoding models for four PFN types. Previous studies have developed a vector encoding model for PFN steady-state responses. Here we extended that approach to allow for encoding of dynamically changing multi-sensory input. To validate our model, we used it to predict responses to complex multi-sensory inputs not used to fit the model. Overall, we found that our model predicted responses to these stimuli well, although a slow decay was present in the real data that was not well-captured by our model, and responses to optic flow in our test measurements were weaker than those in the initial set of experiments to which we fit our model.

We then combined the output of our dynamic encoding model with a nonlinear observability analysis to determine whether this representation is sufficient to compute ambient wind direction. We found that the PFN representation was sufficient to decode wind direction to within ±45° (*i.e.,* quadrant accuracy), when combined with natural “anemometric” flight maneuvers. Surprisingly, we found that few single PFN types had a strong impact on the observability of wind direction. This is because most of the variables needed to decode wind direction are redundantly represented across multiple PFN types. Our models make specific predictions about behavioral deficits that should arise when particular neuron types are silenced. Future experiments in which different PFN types are silenced during flight should allow us to directly test these predictions.

We also constructed and trained artificial neural networks to gain insight into how the fly nervous system might decode ambient wind direction from PFN representations. We found that a simple feedforward ANN with a single hidden layer could decode wind direction to within +/-45°, although the performance of this ANN was slightly worse than the performance of real flies. For both the ANN, and real flies, we found that the ratio of windspeed to groundspeed was a key determinant of success. When this ratio is greater than one, airflow direction can be used as a proxy for ambient wind direction and both the ANN and freely flying flies show superior performance. Real flies show a characteristic deceleration in response to optogenetic odor presentation that increases this ratio, and may contribute to their decoding ability. However, we also found that both flies and ANNs can decode wind direction when this ratio is less than one, more accurately than would be expected from a proxy like airflow direction. A model involving recurrent connections—which are prevalent throughout the Central Complex (*32*)—might improve the performance of our ANN. A more biologically realistic model of wind decoding might also use different temporal filters to represent self-motion history, rather than two distinct inputs representing different time points.

In this study, we combined extensive imaging with computational approaches to show that the multi-sensory representation of self-motion in PFNs is sufficient for a flying fly to decode ambient wind direction when combined with active maneuvers. Several pieces of evidence point to the CX playing a role in wind orientation in walking flies. Stimulation of several tangential neurons and local neurons of the fan-shaped body can drive upwind running (*87*, *88*). Developmental manipulation of the CX— by knocking down the early-expressed RNA binding protein Imp in the type II stem cells that generate the columnar structure of the CX— completely abolishes upwind walking in response to odor (*89*). Local neurons within the fan-shaped body exhibit odor-evoked signals that correlate with the animal’s goal direction both during odor (*90*, *91*) and when returning to an odor plume (*92*). Our data and modeling are consistent with the idea that the fan-shaped body might build an abstract representation of wind direction that allows flies to navigate towards odor sources both in walking and in flight. Experiments manipulating PFN activity in freely flying flies, and ideally, measuring activity under conditions that better simulate free flight, will be required to directly test this hypothesis.

## Methods

### Fly husbandry

Flies for imaging (*Drosophila melanogaster*) were reared and maintained on standard cornmeal-agar media at 25°C, on a 12-hour light cycle (9 AM lights on, or 1 PM lights on for flies imaged in the afternoon). Only mated adult females aged 6-12 days post-eclosion at the time of imaging were used in imaging experiments; these were sorted into culture vials using CO_2_ anesthesia at least five days prior to imaging.

Flies used in free-flight recordings in the wind tunnels were maintained on a 16:8 hour light/dark cycle at 25°C. For optogenetic lines, culture media for parental crosses and for experimental F1 flies was enriched with 400 μL of a 40 mM all-*trans*-retinal solution in ethanol (R2500 from Sigma Aldrich) dissolved in a 15% sucrose solution. Flies were collected at least 48 hours prior to experiments. All free-flight flies were 3-7 days post-eclosion at the time of recording.

### Fly stocks

| Stock Name | Stock ID | Source |
| --- | --- | --- |
| SS00078/([CyO];[TM6b]) | BID#75854 | Derived from hemidriviers |
| SS52244/[CyO] | BID#86596 | BDSC |
| SS02255/[TM6b] | BID#75923 | BDSC |
| SS47384 | BID#86590 | BDSC |
| SS46525 | BID#86589 | BDSC |
| 20xUAS-IVS-jGCaMP7f | BID#80906 | BDSC |
| UAS-tdTomato | BID#36327 | BDSC |
| NorpA-;; | BID#9047 | Stavropoulos Lab |
| ;orco-Gal4; | BID#26818 | Vosshall Lab |
| ;;UAS-Chrimson | BID#55136 | BDSC |

Genotypes used in each figure were as follows:

|  |  |
| --- | --- |
| Figures 1D-K, 2D,F,G, 4B-D,F,H, 5C-F<br>Supp. 1A-I, 3A, 4D,G-J , 5 | SS00078-spGal4 > UAS-GCaMP7f,UAS-tdTomato |
| Figures 2A-G, 3C, 4B,E,H, 5C-F<br>Supp. 2A, 3A, 4A,B,F, 5 | SS52244-spGal4 > UAS-GCaMP7f,UAS-tdTomato |
| Figures 3A,B, 5C-E<br>Supp. 3A, 4C,E | SS02255-spGal4 > UAS-GCaMP7f,UAS-tdTomato |
| Figures 3D-F<br>Supp. 3B-D | SS47384-spGal4 > UAS-GCaMP7f,UAS-tdTomato |
| Figures 3G-I<br>Supp. 3E,F | SS46525-spGal4 > UAS-GCaMP7f,UAS-tdTomato |
| Figure 8, Supp. 9 | NorpA/+;orco-Gal4/+;UAS-CsChrimson/+ |

### Imaging Preparation

Flies were briefly cold-anesthetized before mounting on a custom-made imaging chamber. For open-loop experiments, the chamber was a thin steel foil molded into a dish and affixed to a 3D-printed frame (Clear Resin) with a hexagonal hole for the fly’s head and thorax. For all flight experiments, a separate chamber was 3D-printed with opaque Gray Resin V4 (FormLabs) on a Form3+ 3D printer. While the fly was anesthetized, the two most distal segments of all six legs were removed and the fly was glued to the chamber using UV-cured glue applied around the head and to two points on the thorax. The head and thorax were angled such that the posterior head cuticle was easily accessible through the chamber window. The proboscis was then gently extended using forceps and fixed at the base using the UV glue, so that the labella were free to open and the fly could be fed and watered while mounted. The fly was allowed to recover for at least 30 minutes before the cuticle was dissected away from the head to expose the posterior side of the brain in a bath of artificial hemolymph (AHL: 103 mM NaCl, 3 mM KCl, 5 mM TES, 8 mM trehalose dihydrate, 10 mM glucose, 26 mM NaHCO_3_, 1 mM NaH_2_PO_4_H_2_O, 1.5 mM CaCl_2_2H_2_O, and 4 mM MgCl_2_6H_2_O, pH 7.1-7.4, osmolarity 270-274 mOsm), bubbled with carbogen (5% CO_2_, 95% O_2_).

### Calcium Imaging

To image, each paired structure (protocerebral bridge halves or noduli) was centered in the field of view. The imaging area for the protocerebral bridge ranged from 137.73 x 68.86 μm to 164.10 x 123.07 μm, and the imaging area for the noduli ranged from 74.11 x 49.41 μm to 84.67 x 56.45 μm. Volumes were taken in depth to record the full structures, at a range of 20-32 μm thick and at a rate of 2.6-4.3 volumes per second. Genetically expressed fluorophores (GCaMP7f (*93*) and tdTomato) were simultaneously excited with a pulsed 2-photon laser (Spectra-Physics MaiTai HP) with wavelength 920 nm. Emitted photons were separated by wavelength using a dichroic mirror and bandpass filters (Semrock, FF01-525/50-32 for green and FF01-607/70-32 for red) and collected by GaAsP PMTs. For all open-loop experiments, we used a 20x Olympus water-immersion objective (XLUMPLFLN) with working distance 2.0 mm. For all simultaneous behavior and imaging experiments (Fig. 5 and Supp. Fig. 5), we used a 40x Nikon water-immersion objective (CFI APO NIR) with working distance 3.5 mm. Intensity at the sample was 20-55 mW using the 20x objective, and 10-35 mW using the 40x objective.

### Sensory Stimuli

To present airflow to the fly, we used a rotary union apparatus (*90*) to deliver charcoal-filtered air at a flow rate of 0.2, 0.45 or 0.6 L/min, corresponding to 20, 45, and 60 cm/s. Flow rate was set by a mass flow controller (Aalborg, GFC17 0-2 L/min). Airspeeds were confirmed using a filament anemometer from Dantec Dynamics (model: 9055P0161).

Translational optic flow stimuli were generated using custom Python code and projected onto a 2.75-mm-thick flat, plastic screen 6.5 cm in front of the fly using a DLP 4500EVM and a single flat mirror (3 mm thick) angled 45° relative to the screen. The stimulus filled approximately 90 azimuthal degrees and 60 vertical degrees of the fly’s frontal visual field. The projector framerate was consistently 178-180 Hz, measured by photodiode (ThorLabs, PBM42), and the visual experience was updated at a rate of 50 Hz. To reduce imaging confounds from the blue visual stimulus, we installed a narrow bandpass filter permitting only blue light (Semrock BrightLine FF01-470/28-25) and a neutral density absorptive filter (Thorlabs, NE04B) into the path of the light coming from the projector.

Using the Panda3D package (*94*), blue virtual spheres subtending up to 23° (at the sphere’s closest virtual distance to the fly) of the fly’s visual field randomly populated a black virtual space around the fly. During the optic flow stimulus, the fly’s virtual position in the space was moved in the direction specified without changing its heading direction. While spheres were generated in a large virtual space, for the duration of each trial only those spheres between 5 and 60 units virtual distance in depth from the fly at each timepoint were displayed. To match the optic flow rates to the airflow speeds, we calculate that one virtual distance unit equals 3 cm in real space.

For slip experiments, the optic flow direction was simultaneously slipped at the same speed as the airflow. However, the optic flow stimulus code for the slip data did not utilize a 3D virtual space populated with spheres; instead, it merely produced a dot expansion stimulus from an origin point on-screen. When slipped, this generated a change in yaw in addition to a change in translational travel direction, as it slipped the origin of expansion to the right or left. This artifact was corrected in the 3D virtual space code used in all other experiments reported in this manuscript. In the same experiments, the airflow stimuli were “slipped” 45° to the fly’s right or left at a rate of approximately 93.75°/s, starting at the beginning of the 11th second of the 20-second stimulus.

For imaging experiments with simulated turns in wind, visual and mechanosensory experience of flight through wind was computed using model predictive control with code available in the *pybounds* package (*10*) (further described below). All six of the simulated trajectories’ flight speed began at 20 cm/s and underwent deceleration to approximately half-speed. Two of the trajectories exhibited no turn, while two made a 90° change in course direction to the left and two made the same turn to the right over 4.5 s. For each pair of trajectories with similar course directions (no turn, left turn, and right turn), one was simulated with lower control than the other, governing the influence of wind on the resulting trajectory and the multisensory experience. For all six trajectories, wind was 45 cm/s from 45° relative to the fly’s initial course direction.

For simultaneous recording of behavior and calcium activity, the optic flow stimulus populated only the lower half/third of the region of the fly’s field of view subtended by the visual stimulus screen, producing a horizon-like visual effect. The upper half of the visual stimulus screen contained a “sun dot”, a larger sphere (subtending approximately 10° of the fly’s field of view) that moved laterally with changes in heading but was always the same size on the screen as it maintained a constant 40-unit virtual distance from the fly.

### Flight Behavior Recording

For simultaneous recording of flight behavior and neuronal calcium, the custom imaging chamber was modified to permit the fly’s wings to flap in flight and to mount two lavalier microphones (Audio-Technica, CAT#AT899) posterior to and on either side of the fly.

After mounting and after cuticle dissection to expose the brain, the flies were fed a small amount of sucrose diluted in water, via a Kimwipe soaked in the solution touched to their proboscis, to supply them with energy. Flies that did not drink the solution or were otherwise lethargic were not imaged.

Once the flies were mounted beneath the 2-photon objective, the microphones were placed in their holders below and behind the fly, their distances relative to the fly were balanced, and the threshold signal for flight behavior detection was adjusted during the first 2-10 trials during which flight behavior (wing flapping) was encouraged. Flight was elicited by briefly puffing air from behind the fly through a tube fed through the light-protective box around the imaging setup, such that the flies could be encouraged to fly/flap wings while imaging was ongoing. During these initial habituation/testing trials we delivered both frontal airflow and closed-loop optic flow feedback to the fly when it started flying. Flies that did not fly in these first trials were not further imaged, and their data were not included in final analysis.

Behavior and imaging data were recorded in 30-s trials, with one of four randomized sensory feedback conditions triggered by wing flapping as recorded by the microphones. Sensory feedback started when wing flapping was detected and ended when wing flapping ceased. The four conditions included: airflow from the front of the fly (0°), closed-loop optic flow, both frontal airflow and closed-loop optic flow, and neither sensory experience. For all four conditions, a sun dot was presented 5 virtual units to the left or right of center at the beginning of the trial and moved with the fly’s changes in heading (see details in Sensory Stimuli methods).

### Imaging Data Analysis

Imaging volumes were first processed by computing the mean z projection to generate single frames of data through time. A NoRMCorre rigid motion correction (*95*) was then performed on the red (tdTomato) channel and the same shifts were applied to the green (GCaMP) channel.

Following motion correction, ROIs were drawn by hand in ImageJ/FIJI using the tdTomato/red signal channel. For the protocerebral bridge, one region was drawn around each PB half, then each half was subdivided by hand into eight columnar regions of similar area. For noduli recordings, one ROI was drawn around each nodulus (left and right). Each two-dimensional ROI was then converted to a mean pixel intensity value at every timepoint, and the intensity time series of a background region was subtracted from the time series for each ROI to correct both for motion in the z-axis and for signal bleed-through from the visual stimulus.

Depending on the analysis, two separate baselines were used to calculate ΔF/F in each columnar ROI. One baseline, F_0_, was computed as the mean activity in the ROI within the two seconds leading up to the onset of an open-loop stimulus. This was used to measure the change in signal amplitude across the PB hemisphere due to the stimulus. ΔF/F_0_ was then computed as the difference in fluorescence at every timepoint from F_0_, divided by F_0_.

The other baseline, F_5_, was used to visualize bump position within columns over time. F_5_ was calculated as the mean of the lowest five percent of the column ROIs’ intensity values over the course of an entire experiment. As with ΔF/F_0_, ΔF/F_5_ was computed as the difference in fluorescence at every timepoint from F_5_, divided by F_5_. These values were then convolved with a 5x7 (time x column) gaussian filter with circular padding (imgaussfilt, MATLAB). To compute protocerebral bridge bump position at every timepoint, ΔF/F_5_ for all column ROIs at every timepoint were fit to a sinusoid and the phase was recorded as the columnar position of the sinusoid’s maximum.

To compute bump structure, we took the mean fluorescent activity (ΔF/F_5_) across the columns in the first stimulus frame of every trial, with fluorescent maxima aligned to column 5.

### Connectome Data

Connectome data was obtained from the hemibrain connectome (v1.2.1, http://neuprint.janelia.org/, (*39*)). Synaptic weights were calculated by summing the total number of synapses between all neurons the two cell types (for all connections >3 synapses), then dividing by the number of neurons in the PFN population, for a per-PFN-neuron synaptic weight.

### Tethered Flight Behavior Data Analysis

To detect flight behavior and flight turns from microphone data, we recorded audio at a rate of 1000 Hz and normalized this signal to have zero mean every 10 samples. Flight state was detected online if the sum of the absolute value of normalized signal of the past 30 microphone samples was above threshold for the fly for either microphone. Turns were detected by comparing the envelopes of the signal from the right and left microphones every 10 samples.

Flight behavior was analyzed post-hoc by filtering the normalized microphone data using a bandpass butterworth filter passing harmonics of wing flapping in the frequency range 320-500 Hz, filtering out signal in the band 80-180 Hz (range of noise at and below the frequency of wing flapping) A full spectrogram of the unfiltered microphone data from initial trials was used to identify the 320-500 Hz band as clean harmonic frequency bands for wing flapping. Flight bouts were extracted using the flight state computed online during experiment, and then matched to both the filtered microphone data and neuronal data frames. Turn bouts shown in Fig. 5E were extracted post-hoc by setting threshold of virtual angular velocity (100°/s) computed during behavior for the closed-loop experience (see above) and selecting the data from -2:4 s around the instant the threshold was exceeded.

### Free-flight Behavior Recording and Analysis

#### Optogenetic Paradigm

A detailed description of the optogenetic wind tunnel system can be found in (*33*). In brief: we developed a real time 3D tracking system for fruit flies flying in a wind tunnel using 12 tracking cameras (acA720, Basler AG, Ahrensburg, Germany) and Braid software (*96*). The real-time 3D position of a fly was streamed using custom ROS nodes written in Python. When a fly entered a pre-defined trigger zone (spanning the middle 30 cm of the axis parallel with the wind, 15 cm wide along the axis perpendicular to the wind, and 15-35 cm of the tunnel’s vertical height), it would randomly be assigned either a flash or sham (no flash) event. Flash events were 675 ms of ∼42 µW mm^-2^ red light, as in (*33*). After a trigger event, a built-in refractory period prevented any other triggering events for 10 seconds.

#### Wind Tunnel System

Data were collected from wind tunnels as described previously (*33*) in two separate collection bouts and in one of two conformations. The first conformation is identical to that found in previous work (*33*), and was used for the highest wind speed (40 cm s^-1^). To generate very low wind speeds, the original system was modified to add extra resistance between the fan array and the laminarizing honeycomb layer by installing two layers of window screening and an additional layer of steel mesh.

The lowest measurable wind speed possible with this conformation was approximately 4 cm s^-1^. Average wind speeds for each of the conditions were recorded using an Alnor Air Velocity Meter (TSI, Alnor Airflow Model TA410). For all windspeeds below 40 cm s^-1^, flight trajectories were recorded in 16-hour bouts, with fans changing speed every 20 minutes, including a still-air control period.

#### Trajectory Inclusion Criteria and Data Analyses

As previously reported (*33*), only trajectories which had at least one full second of tracking after the end of the optogenetic stimulus (or for shams, one full second counting from 675 ms after the fly-triggered an event) were included for analysis. Trajectories were manually inspected for continuous sharp jagged regions characteristic of tracking errors and excluded accordingly.

#### Dynamic encoding model for PFNs

Our dynamic encoding models extend previously established vector models of PFN bump activity (*24*, *31*) to capture the temporal dynamics, speed tuning, and multisensory integration we describe in this manuscript. Bump position in PFNd, PFNa, and PFNv was modeled as a sinusoid that moves in the opposite circular direction to the animal’s heading, as shown and modeled previously (*24*). The amplitude of left and right bumps was then modulated by egocentric airflow and optic flow experience according to the equations described below. Models for each modality were fit separately to unimodal data, then combined and tested against multi-modal data.

To model egocentric encoding of optic flow direction, we fit a cosine direction tuning curve to the the steady-state amplitude of the responses to the five directions of optic flow:

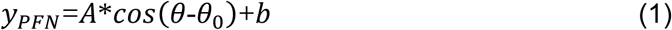

where 𝐴 is the maximal response, 𝜃_0_is the preferred stimulus direction, and 𝑏 is an offset. All fitting was performed using MATLAB’s built-in nonlinear iterative least-squares regression function (nlinfit, MATLAB 2020b).

To model egocentric encoding of airflow direction, we fit Eq. 1 to the average amplitude of the airflow response transients across airflow directions in each PFN type. For PFNa, we augmented this direction tuning curve to have two peaks (as we see in our data and as reported in (*28*)) with a square of cosine term, and with a sum of cosines to produce different peak amplitudes:

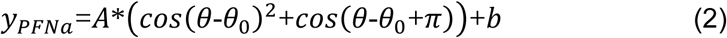

To model airspeed tuning in PFNp_c, we multiplied the direction tuning curve by a saturating exponential function that depended on airspeed and fit this function to the mean amplitude of the airflow response transient to each airflow speed and direction (Supp. Fig. 4A,B).

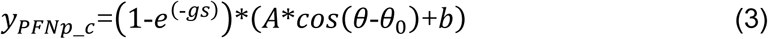

where 𝑠 is airspeed in m/s and 𝑔 is a gain term.

The temporal dynamics of airflow and optic flow responses were described by a single exponential rise (for optic flow) or fall (for airflow). With fixed parameters for direction- and speed-tuning (𝑦_𝑡𝑐_), we fit the remaining parameters to the mean response over time to the stimulus from all directions at all speeds.

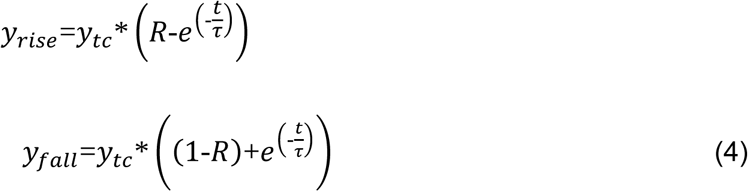

To generate responses to dynamic flight trajectory inputs, we formulated these models as difference equations for each PFN type with the basic form:

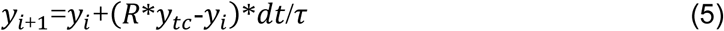

For all simulations, the initial conditions of the PFN responses were assumed to be the steady-state response to the sensory inputs at the first time-step of the simulation.

To capture the multisensory integration strategy in PFNd, we implemented an equal-weight summation of the responses to single-modality stimuli over time. To test the model’s performance, we compared it to actual multisensory responses, which were not used for fitting, by calculating the mean squared error (MSE) between the mean of the multisensory data and the model output over time.

We examined the effect of unequal weighting of the airflow and optic flow response in Supp. Fig. 1F-G using MATLAB’s fitlm function and fitting a two-term linear function to the coherent and divergent PFNd response data:

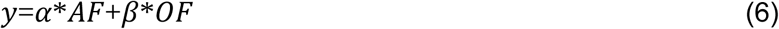

where α is the airflow response weight and β is the optic flow response weight. To compute the additional decay component in Supp. Fig. 4K, we subtracted the equal-weight summation model of coherent activity from the 20-s static coherent response recorded in the stimulus-direction-slip experiment, then fit the parameters of a decay function to the result using MATLAB’s nlinfit:

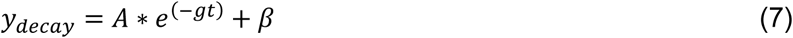

where A is the amplitude, g is a gain term and β is the offset term.

All modeling code was implemented first in MATLAB, then translated into Python for integration with code for model predictive control (*49*) and observability.

#### Wind Direction Observability Analysis

To gauge the capability of the PFN system to estimate the wind direction, we applied an observability analysis. Observability is a concept that describes how difficult a given variable is to estimate, given a set of sensors, or measurements, over time. As applied to our problem, we asked if the ambient wind direction could be inferred from the PFN encodings of heading angle, optic flow direction, and apparent airflow speed and direction.

We employed an established dynamical model of fly flight in the presence of ambient wind that defines the geometric relationship between the state vector 𝑥, which includes the ambient wind vector and a fly’s self-motion variables (heading, ground velocity, etc.), and the corresponding sensor measurement vector 𝑦, which includes heading, optic flow, and apparent airflow (*10*, *11*). We then appended our fit PFN encoding models to the sensor dynamics in this model such that a new measurement vector 𝑦_𝑃𝐹𝑁_ was defined as a function of the original measurements 𝑦_𝑃𝐹𝑁_=𝑓_𝑃𝐹𝑁_(𝑦), where 𝑓_𝑃𝐹𝑁_ defined our fit PFN encoding models. This allowed us to analyze an arbitrary flight trajectory and simulate the corresponding PFN responses over time. We used model predictive control (MPC) to precisely drive our model along desired trajectories (*9*, *49*).

To evaluate the observability of wind direction, we constructed the observability matrix (𝑂) and Fisher information matrix (𝐹) in sliding windows along each flight trajectory as in the *pybounds* python package (*10*). In brief, we empirically computed 𝑂 in each sliding window using our simulation model by applying a small positive and negative perturbation 𝜀 to each initial state variable and simulating the corresponding change in the measurements 𝛥𝑦=𝑦^+^-𝑦^-^ over time:

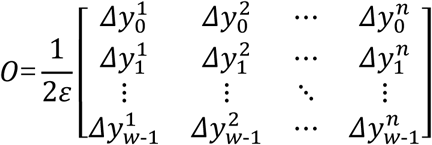

where 𝑛 is the number of states, and 𝑤 is size of the sliding time-window, and 𝜀=10^-5^ (*9*). In plain language, 𝑂 encodes the sensitivity of the model measurements to small change in the initial state and can tell us if each state can be reconstructed from the given measurements.

To compute a quantitative metric for observability, we constructed the Fisher information matrix (𝐹). In general terms, 𝐹 provides information about how much a set of random variables 𝑌 provides about a set of parameters 𝑋. As applied to our dynamical model, 𝑌 is defined by our measurements and 𝑋 is defined by our states. While 𝐹 is typically estimated directly from data (*97*), we construct it from 𝑂 while also considering the presumed noise covariance of measurements. 𝐹 can then be used to compute a quantitative metric for observability given. 𝐹 is defined as

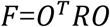

where 𝑅 sets the measurement noise covariance matrix (*9*, *45*).

We set 𝑅=0.1𝐼_𝑘×𝑘_ where 𝐼_𝑘×𝑘_ is the identity matrix of size 𝑘 equal to the number of rows in 𝑂. The Cramér–Rao bound states that the inverse of 𝐹 is a lower bound on an unbiased estimator’s error variance (*98*, *99*). In simple terms, the inverse 𝐹^-1^ provides information about how well a given state could be used in units of variance, e.g. radians squared for the wind direction state. Thus 𝐹^-1^ yields a quantitative metric for observability, where small values correspond to high observability (low error variance) and large values correspond to low observability (high error variance). In practice, 𝐹 was often uninvertible, thus we employed a regularized Chernov inverse

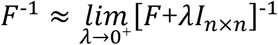

where we set 𝜆=10^-6^. The diagonal elements of 𝐹^-1^ correspond to the minimum error variance (MEV) for each state variable, respectively. We convert this variance into circular variance (MECV) using the following equation:

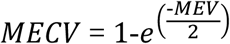

We report the observability of the wind direction throughout as the minimum error circular variance for the wind direction state variable pulled from 𝐹^-1^.

#### Trajectory simulations

In Fig. 6D, all 100-ms simulated trajectories were saccade-like, characterized by a sigmoidal change in course (all in the same direction, towards the fly’s left) and an inverted parabolic change in linear velocity (2.5 m/s^2^ deceleration into the peak of the turn, and equivalent acceleration after the turn) (Supp. Fig. 6A-C). In Fig. 6E, all trajectories shared the same turn amplitude (20°) but varied across deceleration magnitude. In Fig. 6F, all turns were executed with an initial course directionof 90° clockwise rotation away from upwind.

Nearly all trajectories in the paper were simulated using model predictive control input penalties of 0.05 on forward velocity and 0.1 on lateral velocity, producing offsets for trajectory parameters due to wind (*e.g.*, Supp. Fig. 6B,C). The multisensory imaging stimuli in Fig. 4G (used in Fig. 4H, 5F, and Supp. Fig. 5) were produced by running high control MPC trajectory simulation in the presence of a low (5 cm/s) or high (30 cm/s) windspeed, then using the estimated control values from the initial simulation to run the simulation again. Initial simulations with 5 cm/s windspeed produce less controlled secondary simulation trajectories.

In Fig. 6G-H, information (both bump position and half-PB calcium amplitude) from the indicated PFN types were removed from the sensor set used in generating the observability matrix. In Fig. 6I-J, all indicated modality information in all the PFN types was removed from the PFN activity models such that their bump position and calcium intensity outputs would not reflect any of that modality’s inputs.

#### Artificial Neural Networks

To construct neural networks capable of estimating wind direction from PFN activity patterns, we designed and trained a set of feedforward artificial neural networks (ANNs). In these ANNs, an input unit corresponded to a PB column of each PFN type at a particular time in a simulated trajectory, with each unit representing an instantaneous PFN column measurement or one 12 ms before. The output layer is a series of 8 units that we trained to produce a sinusoid whose position represents the allocentric wind direction.

ANNs were Sequential models constructed using the *keras* package (*100*). ANNs varied in the number of hidden layers (1 or 2), the number of units per hidden layer (18 or 36), and the number of training epochs (500 or 1000). The three ANNs chosen for this manuscript were selected based on their small numbers of trainable parameters (<6k) and layers (1 or 2) and on their performance on their training dataset, including comparing loss to validation loss values for convergence indicating minimal overfitting. Two models had two hidden layers of 36 and then 18 units, and were trained for 500 and 1000 epochs respectively. One model had 18 units in one hidden layer during training, which was reduced to 13 units after training following weight dropout testing.

The input layer for the three ANNs consists of 128 units: 16 units per PFN type (PFNd, PFNv, PFNp_c, and PFNa) corresponding to 8 columns in each half of the protocerebral bridge, taken at two timepoints in the course of a simulated trajectory, 12 ms apart. One timepoint is ‘historical’, *i.e.*, the wind direction at t=0 is being estimated using PFN activity at t=0 and at t=-12 ms. We used ReLU activation from the input layer to the hidden layer and between hidden layers, and a linear activation from the last hidden layer to the output layer. Loss was computed as the mean squared error, and the learning rates were optimized using Adaptive Moment Estimation (‘adam’).

The training and test datasets were of modeled PFN activity during 50-ms long simulated trajectories. Possible trajectory parameters were drawn from the following values:

Initial groundspeed = [0.15, 0.4, 0.65, 0.9] m/s

Deceleration = 50 m/s^2^

Turn amplitudes = [-2.639, -1.885 -1.131, -0.377, 0.377, 1.131, 1.885, 2.639] rad

Initial course direction = [-3.122, -2.094, -1.047, -0.698, 0.02, 0.698, 1.047, 2.094] rad

Windspeed = [0, 0.15, 0.3, 0.45, 0.6] m/s

Wind directions = [-3.102, -2.403, -1.705, -1.007, 0.389, 1.087, 1.785, 2.483] rad

Altitude = [0.3, 1.0, 2.0] m

The test dataset used initial parameters pseudorandomly (python *random* module, seed = 29464) selected from [0:1] m (or m/s) or [0:2π] rad, as appropriate for each variable. These trajectories were created with naturalistic heading orientation estimated using the method described below.

After training, the smallest ANN (1,926 trainable parameters, 1 hidden layer of 18 units) performed acceptably well to move forward with experimentation. Necessity testing was done to eliminate units that did not contribute to the wind direction estimate. These had a learned weight parameter near 0 ([-0.3:0.3]) for all input connections. Five hidden units were eliminated using this criterion, resulting in an ANN with 128 input units, one hidden layer with 13 units, and 8 output units.

Because the ANNs invariably produced sinusoidal outputs, all analysis of ANN performance was done by comparing the “columnar” position of the peak of the sinusoid in the output to where the peak lands in the sinusoidal representation of the actual wind direction. To gauge ANN performance without the contributions of individual PFN types, the weights on the input units corresponding to each PFN type were selectively replaced with zeroes. To gauge ANN performance without the contributions of individual modalities (e.g., heading, airflow, optic flow), the weights were left intact but the PFN activity input was adjusted so that encoding of each modality was absent, as described in the Observability methods.

#### Heading Estimation for Measured and Simulated Trajectories

To estimate naturalistic body yaw orientation (heading) for both measured translational flight trajectories and simulated trajectories, we used an artificial neural network approach similar to that described in detail elsewhere (*53*). In brief, we used a fully connected feedforward neural network that was trained on an existing dataset of flying *Drosophila* performing olfactory navigation under multiple wind conditions (*3*, *53*). The training data had undergone randomized rotational shifts to produce experiences in multiple wind direction conditions, but did not also include the original dataset (which only referenced a single wind direction). This difference did not materially change the performance for the neuronal-circuit-inspired ANNs in Fig. 7 and 8 (Fig. 7H and Supp. Fig. 7B). These data included both the translational trajectory data (position and velocity over time) and body yaw orientation over time. The inputs to the heading estimator network consisted of a short time history (a total of 4 consecutive measurements) of ground velocity, air velocity, and estimated thrust vectors in the x-y plane. The network was trained to predict the sine and cosine of the observed body yaw orientation angle. During inference, the time series of these sine and cosine terms was low pass filtered before being converted into a time series body yaw orientation angles.

### Statistical Testing

All distribution comparisons for the ANNs and the real fly data in Figures 7, 8, and Supplemental Figure 9 were tested for significance using Kuiper’s two-sample test, the circular analog of the Kolmogorov-Smirnov test for significant difference between two samples, or Rayleigh’s test for non-uniformity of a circular distribution to compare to chance where indicated. Test code functions were sourced from the *pycircstat* Python module by user philippberens on Github. Due to the very large sample size of the datasets, running a single test on any comparison produced extremely small p-values. To gauge the relative significance of the differences between distributions, data were resampled in subsets of n=501 and the Kuiper test run 10,000 times. Each resampling was performed without replacement, to maximize the chances of sampling most of if not all of the full dataset across the repeated tests. The mean of the 10,000 resulting p-values are reported.

## Data and code availability

All data collected in this manuscript will be made publicly available on Zenodo upon publication. All code (encoding model and observability) as well as CAD files for fly holders is available at Github at https://github.com/nagellab/Mayetal2026.

## Acknowledgements

The authors thank David Schoppik, David Schneider, and Dora Angelaki for input on the project.

## Funding

Leon Levy Scholarship in Neuroscience (C.E.M.)

National Science Foundation Postdoctoral Fellowship in Biology (B.C.) National Institutes of Health grant R01NS136988 (F.v.B.)

National Institutes of Health grant R01DC107979 (K.I.N.) National Institutes of Health grant R01NS127129 (K.I.N.)

## Author Contributions

Conceptualization: C.E.M., B.C., F.v.B., K.I.N.

Methodology: C.E.M., B.C., N.M., F.v.B., K.I.N.

Investigation: All experiments were performed by C.E.M. except the acquisition of real-fly flight behavior datasets by S.D.S and A.L.

Visualization: C.E.M. Supervision: F.v.B., K.I.N.

Writing— original draft: C.E.M., K.I.N.

Writing—review and editing: C.E.M., B.C., F.v.B., K.I.N.

## Competing Interests

All authors declare they have no competing interests.

## Data and materials availability

All data collected in this manuscript will be made publicly available on Zenodo upon publication. All code (encoding model and observability) as well as CAD files for fly holders will be available at Github at https://github.com/nagellab/Mayetal2026.

---

**Supplement 1 related to Figure 1.**
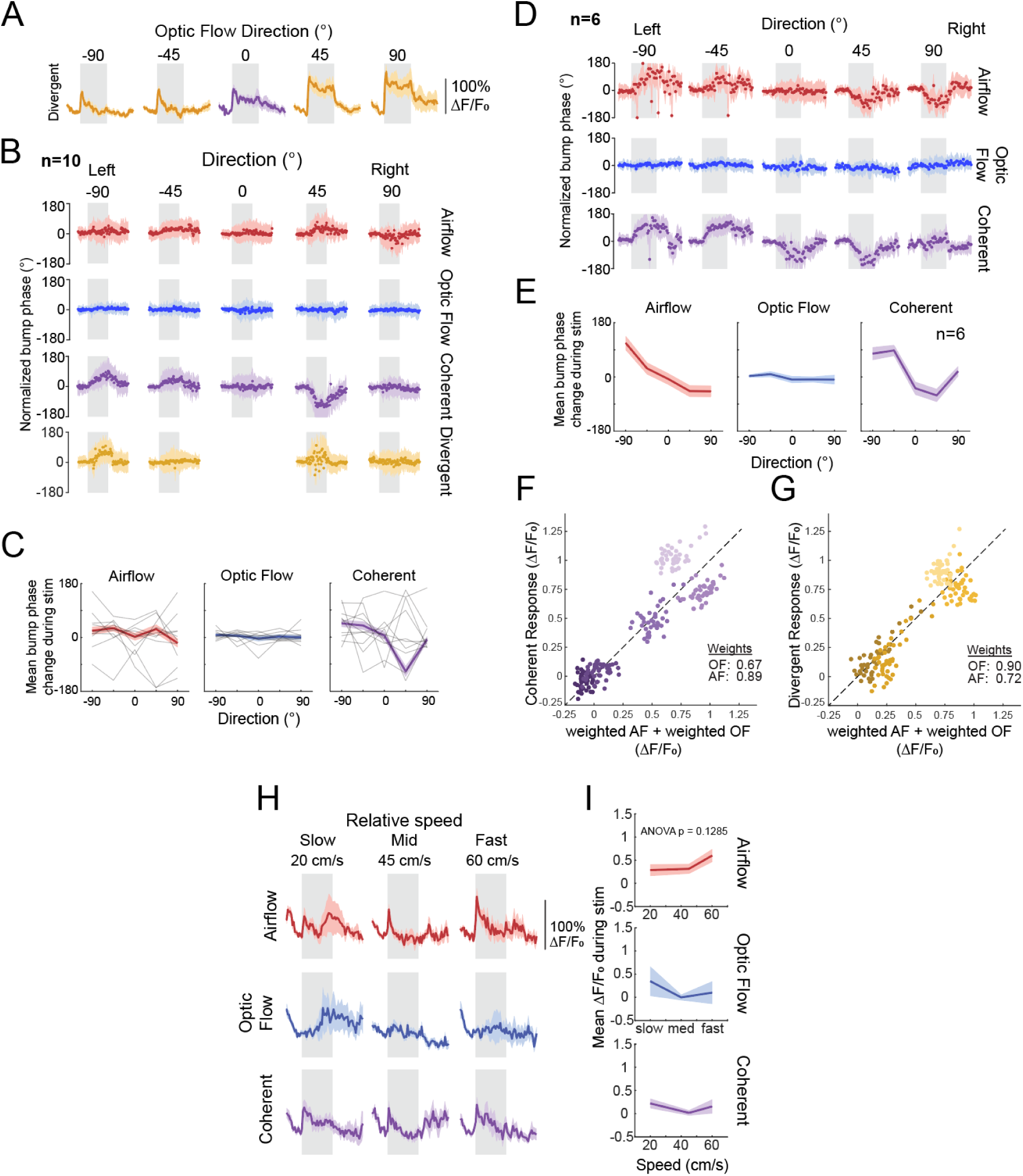
Additional data on PFNd responses. A. Calcium activity dynamics in PFNd PB halves to divergent multisensory stimuli. Airflow is from 0° for all stimuli; translational optic flow is from the directions indicated. The response to coherent airflow and optic flow from 0° is reproduced from Fig. 1F. Gray areas indicate stimulus period. Colored traces represent mean across flies ±SEM (N=10 flies, 2-4 trials per fly, same data as in Fig. 1F). B. Bump phase across PB columns over time, normalized to phase immediately before stimulus onset for all data. Negative directions indicate stimuli coming from fly’s egocentric left. Data are means of means across flies and trials; light traces are circular means across trials per fly; bold traces and color shading represent circular mean across flies ± circular standard deviation. n=10 flies, 2-4 trials per fly, same data as in Fig. 1F. C. Mean change in bump phase across PB columns relative to phase immediately before stimulus onset for all data. Negative directions indicate stimuli coming from fly’s egocentric left. Light traces are circular means across trials per fly; bold traces are circular means; shading is ±circular standard deviation. n=10 flies, 2-4 trials per fly, same data as in B. D. Bump phase across PB columns over time for a subset of flies (n=6) from B, that exhibit stronger and more consistent bump movement in response to changing airflow direction. E. Mean change in bump phase across PB columns relative to phase immediately before stimulus for subset of flies shown in D. F. Mean activity during the 10-second coherent stimulus (y-axis) versus a weighted sum (see Methods) of the 10-s responses to individual stimuli (x-axis); same data as in Fig. 2G. G. Mean activity during the 10-second divergent stimulus (y-axis) versus a weighted sum (see Methods) of the 10-s responses to individual stimuli (x-axis); same data as in Fig. 2I. H. PFNd calcium activity in response to three stimulus speeds delivered from 0°. Traces represent mean ±SEM across flies for each PB half (N= 6 flies, 2-4 trials per fly). Gray areas indicate 10 s stimulus period. I. Speed tuning curves for PFNd (1^st^ second of the calcium response), same data as in F Shading is ±SEM across flies.

**Supplement 2 related to Figure 2.**
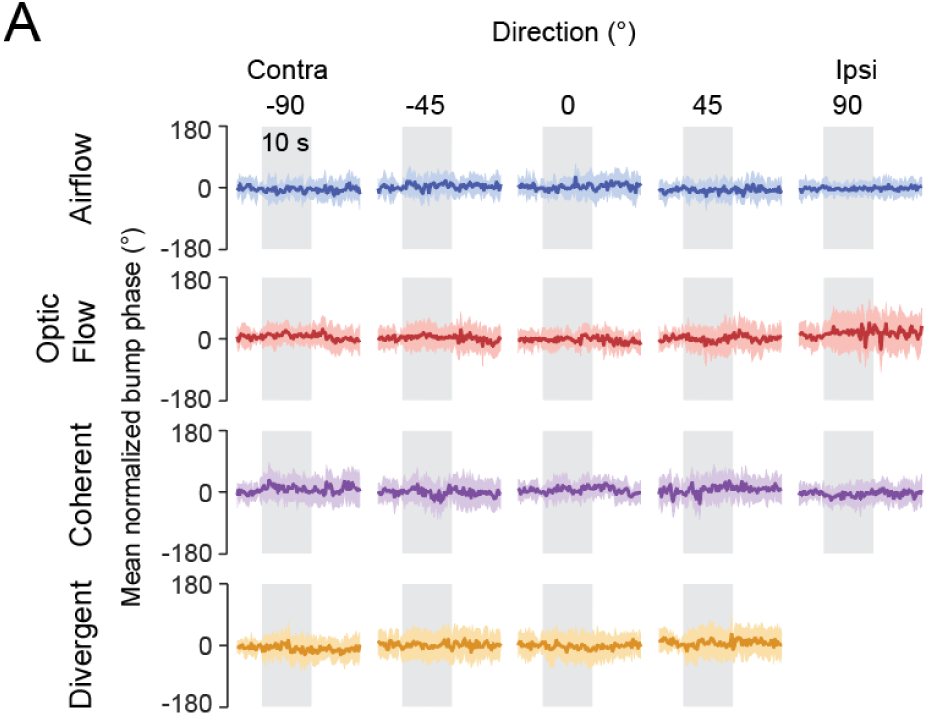
PFNp_c does not exhibit bump movement to any stimulus. A. Mean PFNp_c bump phase across PB columns over time, normalized to phase immediately before stimulus onset for all data. Negative directions indicate stimuli coming from fly’s egocentric left. Traces are circular means; shading is ±circular standard deviation. N=7 flies, 2-4 trials per fly, same dataset as Figure 2E.

**Supplement 3 related to Figure 3.**
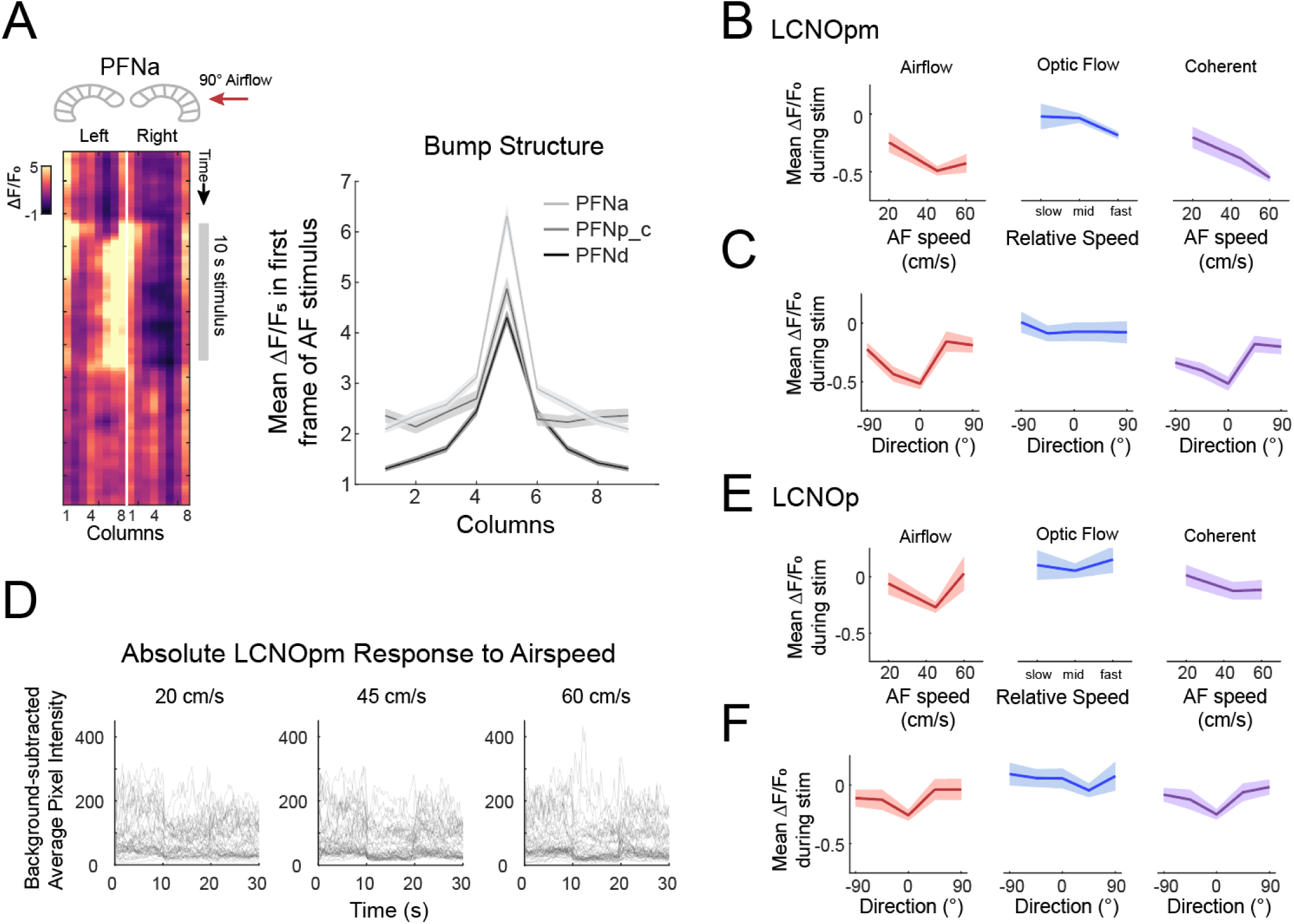
Additional data on airflow tuning in PFNa and LNO neurons. A. (*left*) Example single trial calcium activity in PFNa to 45 cm/s airflow from 90° to the fly’s right. (*right*) Mean bump structure in PFNa (light gray), PFNp_c (dark gray) and PFNd (black). Column 1 is plotted twice (once on either side). Data are means across all trials ±SEM. B. Speed tuning curves (mean ±SEM of the LCNOpm response during the 10-s stimulus period). Same data as Figure 3E. C. Direction tuning curves (mean ±SEM of the LCNOpm response during the 10-s stimulus period). Same data as Figure 3F. D. Mean pixel intensity values of LCNOpm in the NO regions (following mean-z-projection and motion correction; see Methods) minus the mean motion-corrected pixel intensity value of a non-fluorescent background region. Note the nearness of the raw pixel values to zero intensity during the stimulus. Stimulus occurred from 10-20 s. E. Speed tuning curves (mean ±SEM of the LCNOp calcium response during the 10-s stimulus period). Same data as Figure 3H. F. Direction tuning curves (mean ±SEM of the LCNOp calcium response during the 10-s stimulus period). Same data as Figure 3I.

**Supplement 4 related to Figure 4.**
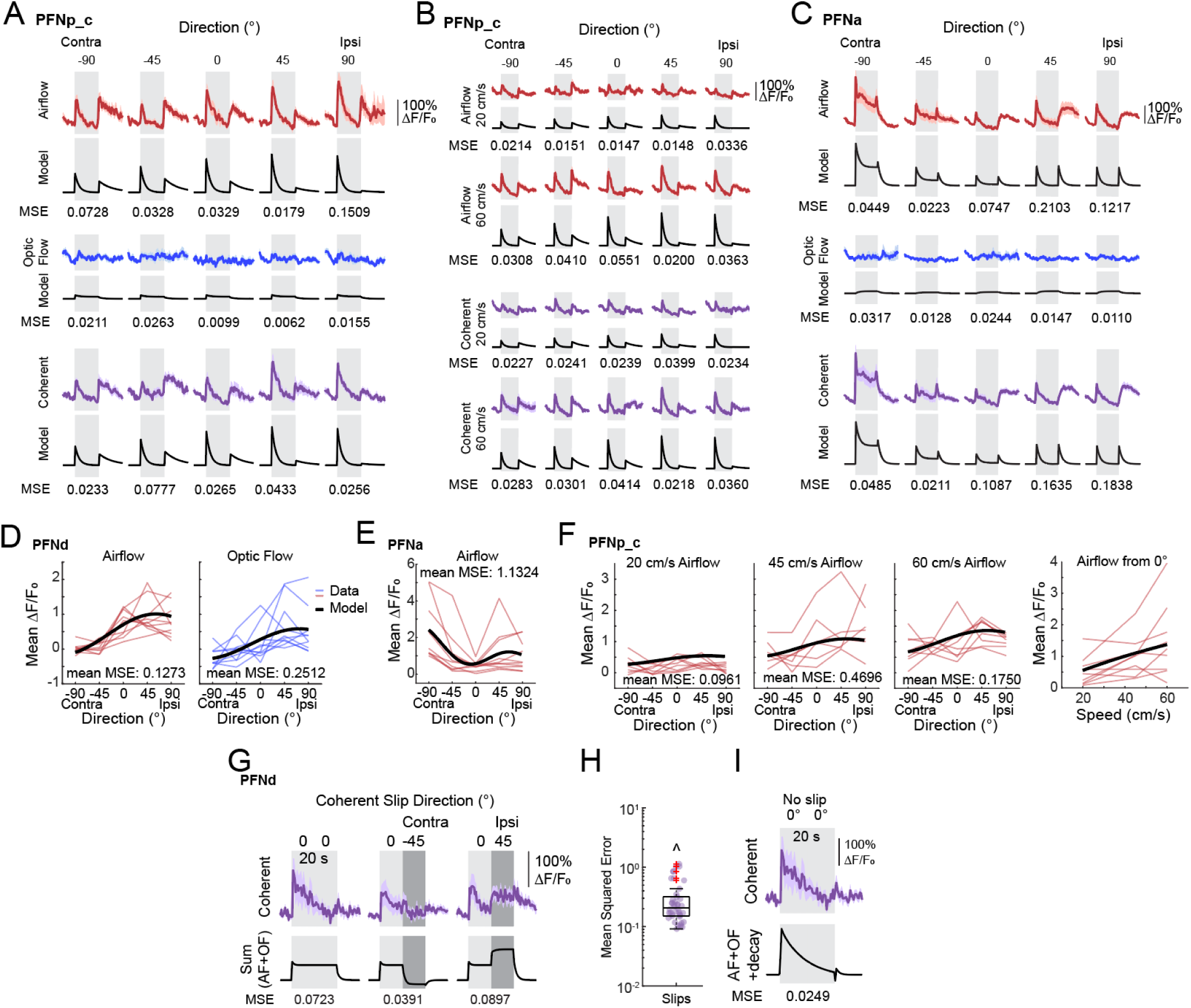
Additional fits and mean squared error for PFN encoding models. A. Real (top, colored traces, reproduced from Fig. 2E) and modeled (black) responses of PFNp_c to 5 directions of stimulus presentation. MSE between model and data shown below each pair of traces. B. Real (top, colored traces, N = 8 flies, 2-4 trials per fly) and modeled (black) responses of PFNp_c to 5 directions of stimulus presentation at two speeds (20 cm/s and 60 cm/s). Colored traces are mean ±SEM, and black traces are the model outputs for each stimulus condition. MSE between model and data shown below each pair of traces. C. Real (top, colored traces, reproduced from Fig. 3A) and modeled (black) responses of PFNa to 5 directions of stimulus presentation. MSE between model and data shown below each pair of traces. D. PFNd direction tuning curves for individual flies during the first second of the stimulus period (colored traces) with model tuning curve overlaid in black. E. Same as D, for PFNa airflow direction tuning. F. Same as D, for PFNp_c airflow direction and airspeed tuning. G. Real (*top*, colored traces) and predicted model (black) responses to multimodal “slip” stimuli. In the left panel, the coherent stimuli are presented statically from 0° during the entire 20-sec stimulus period. In the middle panel, the coherent stimuli are presented from 0° for 10 sec, and then slipped sideways to present from contralateral 45° for 10 sec. In the right panel, the coherent stimuli are presented from 0° for 10 sec, and then slipped to ipsilateral 45° for 10 sec. MSE between model and data shown below each pair of traces. (Data from n=6 flies, 2-4 trials per fly.) H. Box and dot plots of MSEs for PFNd mean response to slip stimuli in G. I. MSE of equal-weight sum of single-modality responses plus a fitted additional decay function (see Methods) vs. coherent data from slip experiment (same data as G).

**Supplemental Figure 5, related to Figure 5.**
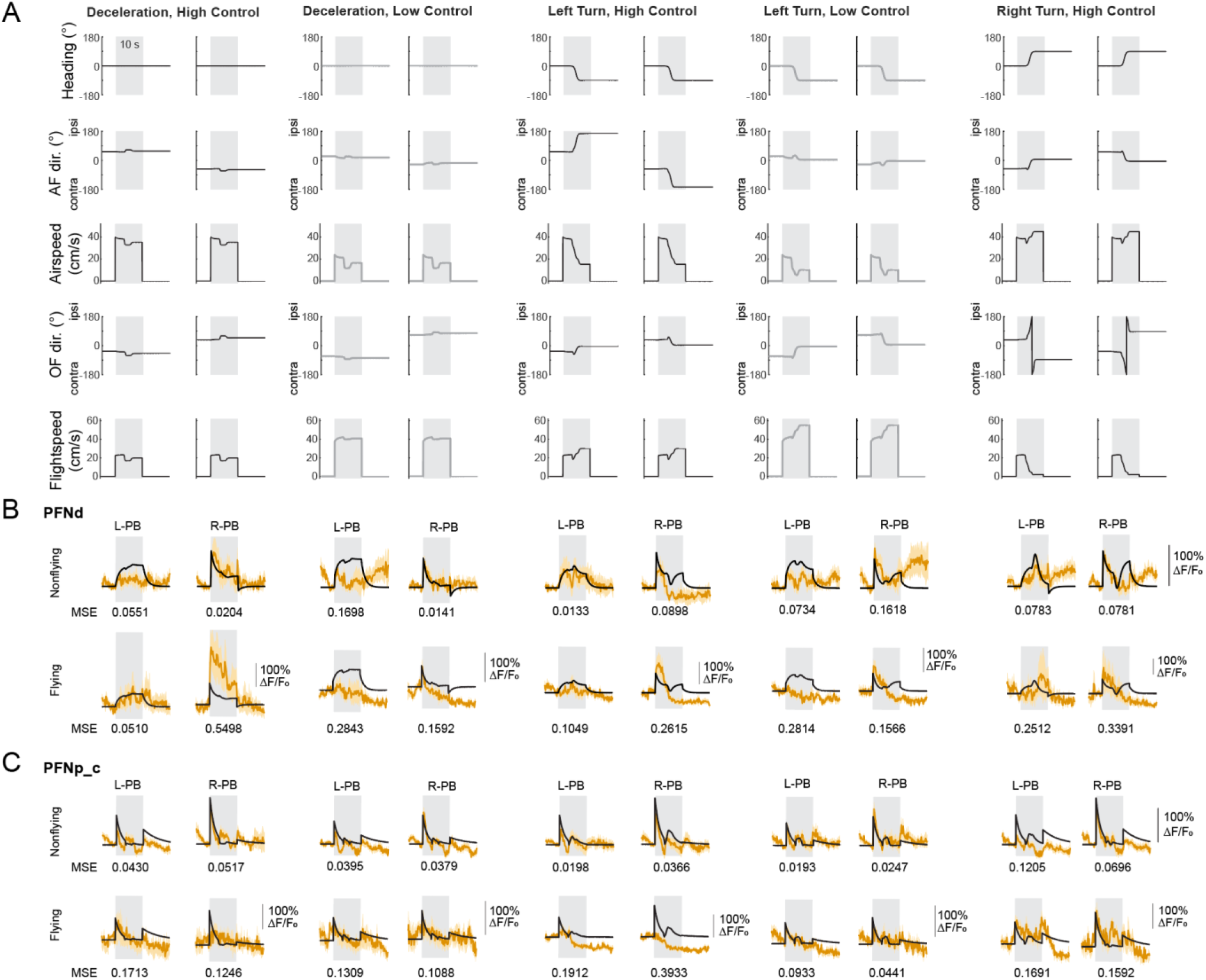
In-flight PFN responses to dynamic stimuli simulating flight through wind. A. All stimulus variables drawn from five dynamic flight-through-wind experiences, with or without a simulated turn, and with either low or high control of the trajectory. Egocentric airflow and optic flow directions are reported as ipsilateral (positive values) or contralateral, matching the PB half response traces below in B and C. B. PFNd responses as mean ± SEM ΔF/F_0_ within the left or right PB in nonflying (top) or flying (bottom) flies to the experience flight through wind with or without a turn, with high or low control. The black traces are encoding model predictions. Each nonflying data trace includes 3-6 trials per fly, with n=10 flies. Each flying data trace includes 3-12 bouts of constant flight during the stimulus, from 7 flies. C. Same as B, for PFNp_c. Each nonflying data trace includes 3-6 trials per fly, with n=10 flies. Each flying data trace includes 3-6 bouts of constant flight during the stimulus, from 6 flies.

**Supplement 6 related to Figure 6.**
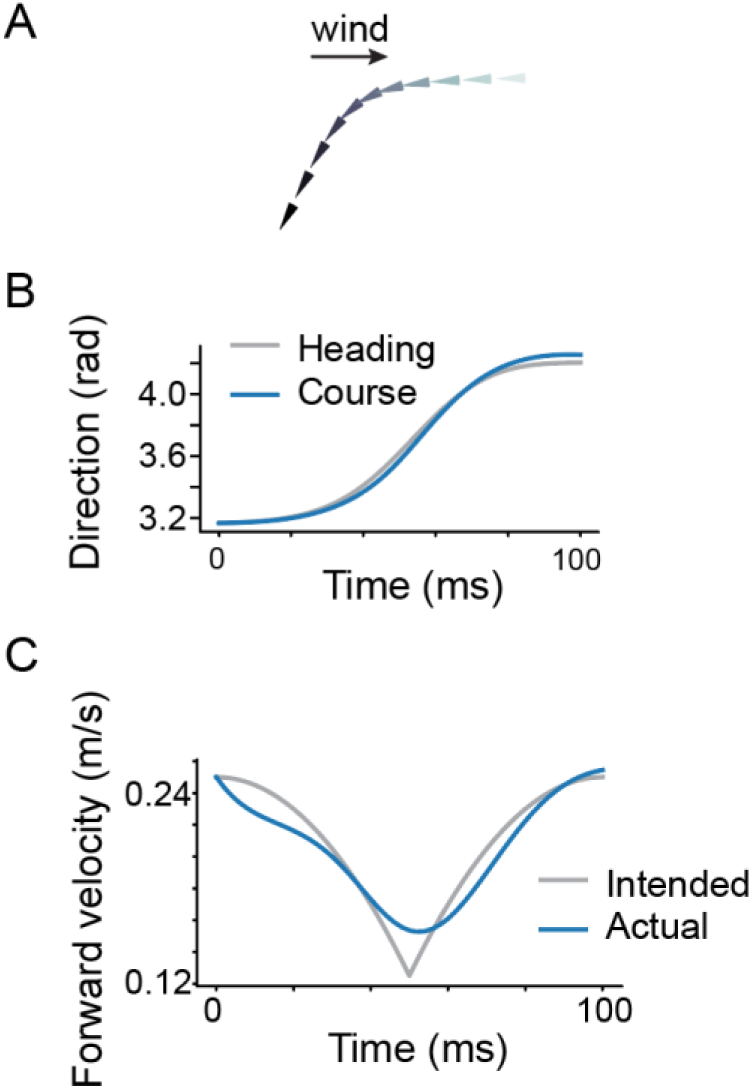
Simulated anemometric turn parameters, and observability analysis using “raw” variable sensors. A. Example simulated saccade-like trajectory with sequential deceleration and acceleration. The arrows represent position and course direction (the pointed direction) at periodic timepoints in the trajectory. B. Heading and travel (course) direction over time for the simulated trajectory in A. C. Forward velocity for the simulated trajectory in A. In gray is the simulated fly’s intended velocity; in blue is the resulting velocity from model predictive control.

**Supplemental Figure 7, related to Figure 7.**
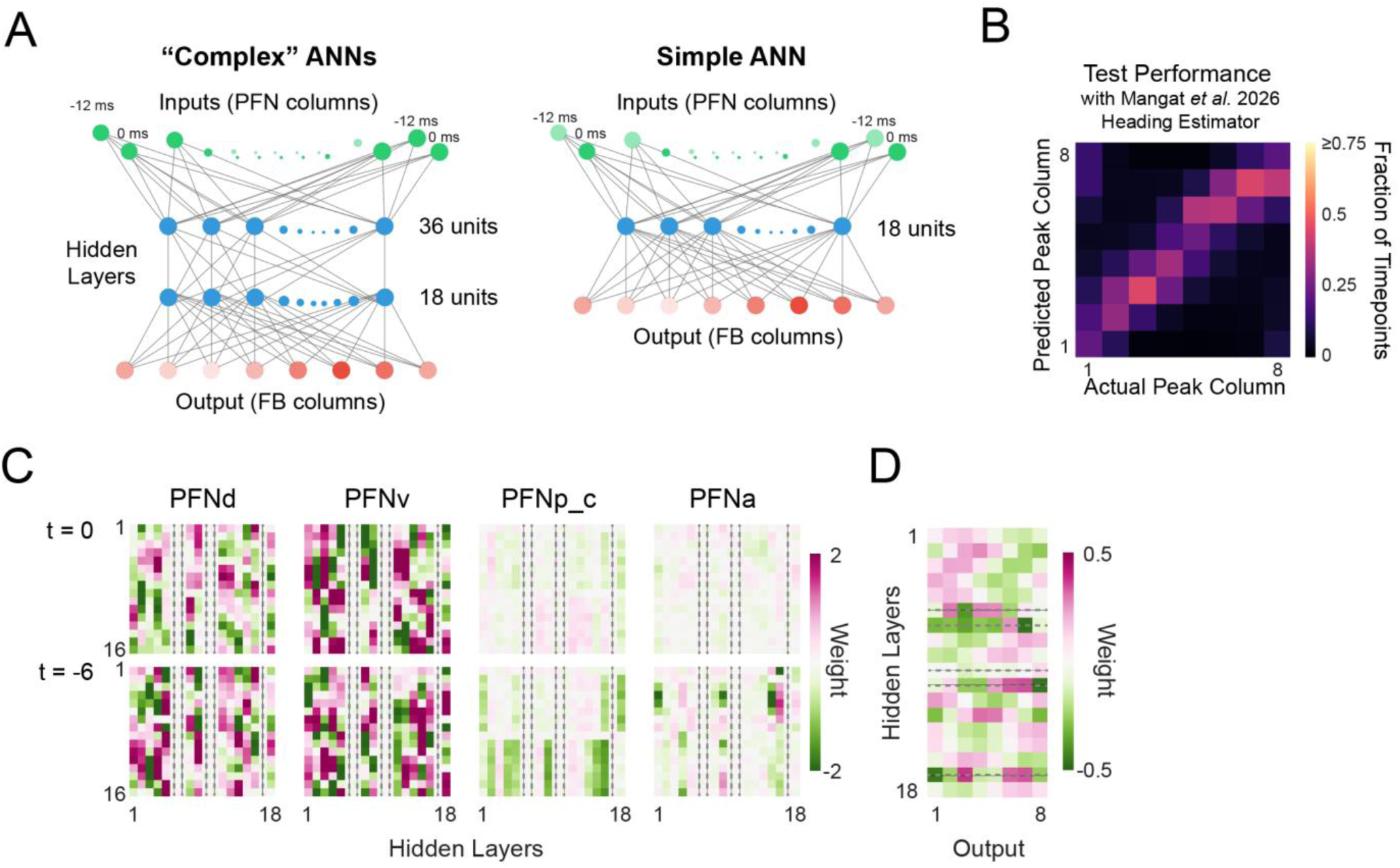
Details about ANN structure and performance. A. Structure comparison between the two more complex ANNs trained in this paper (5,462 parameters, two hidden layers with 36 and 18 units respectively) and the simpler ANN (1,789 parameters, one hidden layer of 18 units (or 13 units after dropout testing)). B. ANN performance on test data with naturalistic headings estimated using the network described in CITE. Compare to Fig. 7H. C. Learned weights from inputs to the hidden layer of the most compact ANN before weight dropout. Eliminated units’ weights are struck-through with dashed lines. D. Learned weights from the hidden layer to the output of the most compact ANN before weight dropout. Eliminated units’ weights are struck-through with dashed lines. Note that these are the only weight sets without sinusoidal structure.

**Supplemental Figure 8, related to Figure 8.**
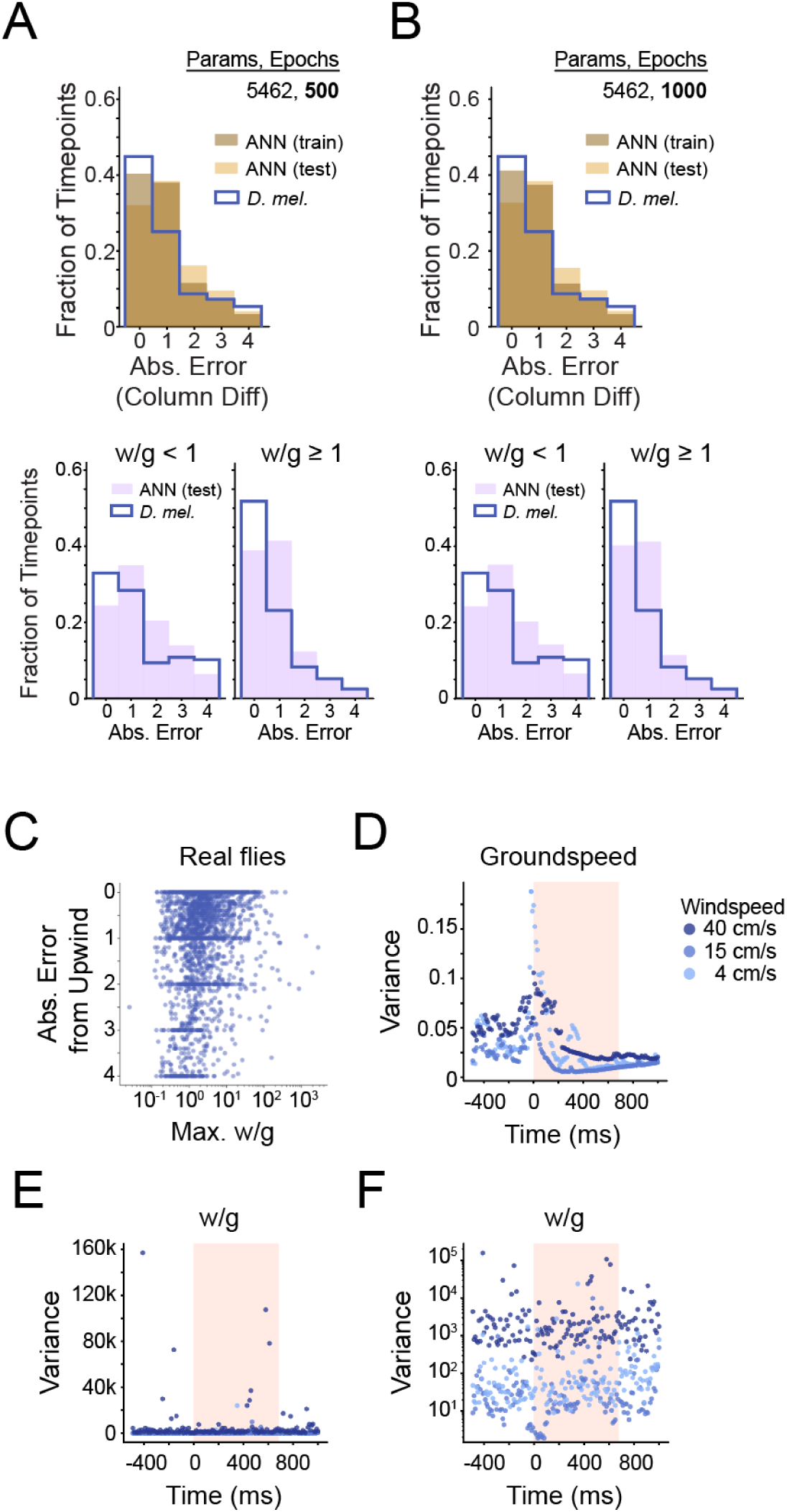
A. (*above*) Comparison between the distribution of wind direction estimation error for a more complex ANN (5,462 parameters, two hidden layers of 36 and 18 units, trained for 500 epochs) and for real flies (same data as Fig. 8F). (*below*) Same comparisons separated into low (*left*) and high (*right*) windspeed to groundspeed ratio. B. Same as A for a second more complex ANN (5,462 parameters, two hidden layers of 36 and 18 units, trained for 1000 epochs). C. Relationship of maximum windspeed-to-groundspeed ratio achieved during anemometric flight maneuvers (deceleration) in real flies (x axis) versus binned course error from upwind (y axis). Course direction bin key in Fig. 8F. D. Variance of real fly groundspeeds over time, colored by ambient windspeed in the wind tunnel. Pink shading indicates duration of fictive odor stimulus by optogenetics. E. Variance of real fly w/g over time, colored by ambient windspeed in the wind tunnel. Pink shading indicates duration of fictive odor stimulus by optogenetics. F. Zoomed-in view of w/g variance in real fly wind tunnel data; same data as F.

## Notes

### Competing Interest Statement

The authors have declared no competing interest.

### Summary of Updates

We have added validation experiments for our PFN encoding model in both non-behaving and flying flies. We also now show how a biologically-inspired feedforward artificial neural network (ANN) can read out ambient wind direction from PFN representations. We show that the ratio of windspeed to groundspeed is a critical parameter governing the performance of both the ANN and of real flies using a newly collected set of free flight behavioral data.

## References

1. W. J. Bell, E. Kramer, Search and anemotactic orientation of cockroaches. J. Insect Physiol. 25, 631–640 (1979).

2. H. Wolf, R. Wehner, Pinpointing food sources: olfactory and anemotactic orientation in desert ants, Cataglyphis fortis. J. Exp. Biol. 203, 857–868 (2000).

3. F. Van Breugel, M. H. Dickinson, Plume-tracking behavior of flying Drosophila emerges from a set of distinct sensory-motor reflexes. Curr. Biol. 24, 274–286 (2014).

4. E. Álvarez-Salvado, A. M. Licata, E. G. Connor, M. K. McHugh, B. M. N. King, N. Stavropoulos, J. D. Victor, J. P. Crimaldi, K. I. Nagel, Elementary sensory-motor transformations underlying olfactory navigation in walking fruit-flies. Elife 7 (2018).

5. J. S. Kennedy, D. Marsh, Pheromone-regulated anemotaxis in flying moths. Science (1979). 184, 999–1001 (1974).

6. E. von Holst, H. Mittelstaedt, Das Reafferenzprinzip - Wechselwirkungen zwischen Zentralnervensystem und Peripherie. Naturwissenschaften 37, 464–476 (1950).

7. K. E. Cullen, L. B. Minor, Semicircular canal afferents similarly encode active and passive head-on-body rotations: implications for the role of vestibular efference. J. Neurosci. 22, RC226 (2002).

8. F. Van Breugel, A nonlinear observability analysis of ambient wind estimation with uncalibrated sensors, inspired by insect neural encoding. Proc. IEEE Conf. Decis. Control 2021-December, 1399–1406 (2021).

9. B. Cellini, B. Boyacioǧlu, F. Van Breugel, Empirical individual state observability. Institute of Electrical and Electronics Engineers Inc. [Preprint] (2023). 10.1109/cdc49753.2023.10383812.

10. B. Cellini, B. Boyacioǧlu, S. David Stupski, F. Van Breugel, Discovering and exploiting active sensing motifs for estimation with empirical observability. bioRxiv, 2024.11.04.621976 (2024).

11. F. Van Breugel, R. Jewell, J. Houle, Active anemosensing hypothesis: how flying insects could estimate ambient wind direction through sensory integration and active movement. J. R. Soc. Interface 19 (2022).

12. H. G. Krapp, R. Hengstenberg, Estimation of self-motion by optic flow processing in single visual interneurons. Nature 384, 463–466 (1996).

13. M. Lappe, F. Bremmer, A. V. Van Den Berg, Perception of self-motion from visual flow. Trends Cogn. Sci. 3, 329–336 (1999).

14. F. Hlavačka, T. Mergner, G. Schweigart, Interaction of vestibular and proprioceptive inputs for human self-motion perception. Neurosci. Lett. 138, 161–164 (1992).

15. J. Dichgans, T. Brandt, Visual-Vestibular Interaction: Effects on Self-Motion Perception and Postural Control. Perception, 755–804 (1978).

16. J. P. Noel, D. E. Angelaki, Cognitive, Systems, and Computational Neurosciences of the Self in Motion. Annu. Rev. Psychol. 73, 103–129 (2022).

17. K. E. Cullen, Vestibular processing during natural self-motion: implications for perception and action. Nat. Rev. Neurosci. 20, 346–363 (2019).

18. N. H. Barmack, H. Shojaku, Vestibular and visual climbing fiber signals evoked in the uvula-nodulus of the rabbit cerebellum by natural stimulation. 10.1152/jn.1995.74.6.2573 74, 2573–2589 (1995).

19. K. Maekawa, J. I. Simpson, Climbing fiber responses evoked in vestibulocerebellum of rabbit from visual system. J. Neurophysiol. 36, 649–666 (1973).

20. K. E. Cullen, Internal models of self-motion: neural computations by the vestibular cerebellum. Trends Neurosci. 46, 986–1002 (2023).

21. A. Chen, G. C. DeAngelis, D. E. Angelaki, Representation of Vestibular and Visual Cues to Self-Motion in Ventral Intraparietal Cortex. Journal of Neuroscience 31, 12036–12052 (2011).

22. Y. Gu, D. E. Angelaki, G. C. DeAngelis, Neural correlates of multisensory cue integration in macaque MSTd. Nat. Neurosci. 11, 1201–1210 (2008).

23. C. R. Fetsch, A. Pouget, G. C. Deangelis, D. E. Angelaki, Neural correlates of reliability-based cue weighting during multisensory integration. Nat. Neurosci. 15, 146–154 (2012).

24. C. Lyu, L. F. Abbott, G. Maimon, Building an allocentric travelling direction signal via vector computation. Nature 601, 92–97 (2022).

25. P. T. Weir, M. H. Dickinson, Functional divisions for visual processing in the central brain of flying Drosophila. Proc. Natl. Acad. Sci. U. S. A. 112, E5523–E5532 (2015).

26. T. Stone, B. Webb, A. Adden, N. Ben Weddig, A. Honkanen, R. Templin, W. Wcislo, L. Scimeca, E. Warrant, S. Heinze, An Anatomically Constrained Model for Path Integration in the Bee Brain. Current Biology 27, 3069–3085.e11 (2017).

27. T. A. Currier, A. M. M. Matheson, K. I. Nagel, Encoding and control of orientation to airflow by a set of Drosophila fan-shaped body neurons. Elife 9, 1–29 (2020).

28. I. G. Ishida, S. Sethi, T. L. Mohren, M. K. Haraguchi, L. F. Abbott, G. Maimon, Neuronal calcium spikes enable vector inversion in the Drosophila brain. Cell 189, 748–764.e25 (2026).

29. L. Polat, T. Harpaz, A. Zaidel, Rats rely on airflow cues for self-motion perception. Current Biology 34, 4248–4260.e5 (2024).

30. S. P. Sane, A. Dieudonné, M. A. Willis, T. L. Daniel, Antennal mechanosensors mediate flight control in moths. Science (1979). 315, 863–866 (2007).

31. J. Lu, A. H. Behbahani, L. Hamburg, E. A. Westeinde, P. M. Dawson, C. Lyu, G. Maimon, M. H. Dickinson, S. Druckmann, R. I. Wilson, Transforming representations of movement from body-to world-centric space. Nature 601, 98–104 (2022).

32. B. K. Hulse, H. Haberkern, R. Franconville, D. B. Turner-Evans, S. Y. Takemura, T. Wolff, M. Noorman, M. Dreher, C. Dan, R. Parekh, A. M. Hermundstad, G. M. Rubin, V. Jayaraman, A connectome of the Drosophila central complex reveals network motifs suitable for flexible navigation and context-dependent action selection. Elife 10, e66039 (2021).

33. S. D. Stupski, F. van Breugel, Wind gates olfaction-driven search states in free flight. Current Biology 34, 4397–4411.e6 (2024).

34. J. S. Kennedy, The Visual Responses of Flying Mosquitoes. Proceedings of the Zoological Society of London A109, 221–242 (1940).

35. J. A. Bender, M. H. Dickinson, A comparison of visual and haltere-mediated feedback in the control of body saccades in Drosophila melanogaster. J. Exp. Biol. 209, 4597–4606 (2006).

36. T. S. Okubo, P. Patella, I. D’Alessandro, R. I. Wilson, A neural network for wind-guided compass navigation. Neuron 107, 924–940 (2020).

37. T. Wolff, M. Eddison, N. Chen, A. Nern, P. Sundaramurthi, D. Sitaraman, G. M. Rubin, Cell type-specific driver lines targeting the Drosophila central complex and their use to investigate neuropeptide expression and sleep regulation. bioRxiv, 2024.10.21.619448 (2024).

38. N. Eckstein, A. S. Bates, A. Champion, M. Du, Y. Yin, P. Schlegel, A. K. Y. Lu, T. Rymer, S. Finley-May, T. Paterson, R. Parekh, S. Dorkenwald, A. Matsliah, S. C. Yu, C. McKellar, A. Sterling, K. Eichler, M. Costa, S. Seung, M. Murthy, V. Hartenstein, G. S. X. E. Jefferis, J. Funke, Neurotransmitter classification from electron microscopy images at synaptic sites in Drosophila melanogaster. Cell 187, 2574–2594.e23 (2024).

39. L. K. Scheffer, C. S. Xu, M. Januszewski, Z. Lu, S. Y. Takemura, K. J. Hayworth, G. B. Huang, K. Shinomiya, J. Maitin-Shepard, S. Berg, J. Clements, P. M. Hubbard, W. T. Katz, L. Umayam, T. Zhao, D. Ackerman, T. Blakely, J. Bogovic, T. Dolafi, D. Kainmueller, T. Kawase, K. A. Khairy, L. Leavitt, P. H. Li, L. Lindsey, N. Neubarth, D. J. Olbris, H. Otsuna, E. T. Trautman, M. Ito, A. S. Bates, J. Goldammer, T. Wolff, R. Svirskas, P. Schlegel, E. R. Neace, C. J. Knecht, C. X. Alvarado, D. A. Bailey, S. Ballinger, J. A. Borycz, B. S. Canino, N. Cheatham, M. Cook, M. Dreher, O. Duclos, B. Eubanks, K. Fairbanks, S. Finley, N. Forknall, A. Francis, G. P. Hopkins, E. M. Joyce, S. Kim, N. A. Kirk, J. Kovalyak, S. A. Lauchie, A. Lohff, C. Maldonado, E. A. Manley, S. McLin, C. Mooney, M. Ndama, O. Ogundeyi, N. Okeoma, C. Ordish, N. Padilla, C. Patrick, T. Paterson, E. E. Phillips, E. M. Phillips, N. Rampally, C. Ribeiro, M. K. Robertson, J. T. Rymer, S. M. Ryan, M. Sammons, A. K. Scott, A. L. Scott, A. Shinomiya, C. Smith, K. Smith, N. L. Smith, M. A. Sobeski, A. Suleiman, J. Swift, S. Takemura, I. Talebi, D. Tarnogorska, E. Tenshaw, T. Tokhi, J. J. Walsh, T. Yang, J. A. Horne, F. Li, R. Parekh, P. K. Rivlin, V. Jayaraman, M. Costa, G. S. X. E. Jefferis, K. Ito, S. Saalfeld, R. George, I. A. Meinertzhagen, G. M. Rubin, H. F. Hess, V. Jain, S. M. Plaza, A connectome and analysis of the adult Drosophila central brain. Elife 9, e57443 (2020).

40. F. T. Muijres, M. J. Elzinga, N. A. Iwasaki, M. H. Dickinson, Body saccades of Drosophila consist of stereotyped banked turns. Journal of Experimental Biology 218, 864–875 (2015).

41. B. Cellini, J. M. Mongeau, Hybrid visual control in fly flight: insights into gaze shift via saccades. Curr. Opin. Insect Sci. 42, 23–31 (2020).

42. B. Schnell, I. G. Ros, M. H. Dickinson, A Descending Neuron Correlated with the Rapid Steering Maneuvers of Flying Drosophila. Current Biology 27, 1200–1205 (2017).

43. I. G. Ros, J. J. Omoto, M. H. Dickinson, Descending control and regulation of spontaneous flight turns in Drosophila. Curr. Biol. 34, 531–540.e5 (2024).

44. N. D. Kathman, J. L. Fox, Representation of Haltere Oscillations and Integration with Visual Inputs in the Fly Central Complex. Journal of Neuroscience 39, 4100–4112 (2019).

45. B. Boyacioǧlu, F. Van Breugel, DUALITY OF STOCHASTIC OBSERVABILITY AND CONSTRUCTABILITY AND LINKS TO FISHER INFORMATION A PREPRINT. (2025).

46. A. K. Singh, J. Hahn, On the use of empirical gramians for controllability and observability analysis. Proceedings of the American Control Conference 1, 140–141 (2005).

47. C. Jauffret, Observability and Fisher information matrix in nonlinear regression. IEEE Trans. Aerosp. Electron. Syst. 43, 756–759 (2007).

48. A. J. Krener, K. Ide, Measures of unobservability. Proceedings of the IEEE Conference on Decision and Control, 6401–6406 (2009).

49. F. Fiedler, B. Karg, L. Lüken, D. Brandner, M. Heinlein, F. Brabender, S. Lucia, do-mpc: Towards FAIR nonlinear and robust model predictive control. Control Eng. Pract. 140 (2023).

50. C. Schilstra, J. H. Van Hateren, Blowfly flight and optic flow. I. Thorax kinematics and flight dynamics. Journal of Experimental Biology 202, 1481–1490 (1999).

51. L. F. Tammero, M. H. Dickinson, The influence of visual landscape on the free flight behavior of the fruit fly Drosophila melanogaster. Journal of Experimental Biology 205, 327–343 (2002).

52. A. P. Lopez, F. van Breugel, “Flies, Data, Robots: Insect-Inspired Flight Strategies for Robust UAV Navigation in GPS Denied Environments,” thesis, United States -- Nevada (2025).

53. N. Mangat, C. E. May, K. I. Nagel, F. van Breugel, Predicting Drosophila Body Orientation from a Translational Trajectory using an Artificial Neural Network. bioRxiv, 2026.03.30.715335 (2026).

54. J. Laurens, H. Meng, D. E. Angelaki, Computation of linear acceleration through an internal model in the macaque cerebellum. Nat. Neurosci. 16, 1701–1708 (2013).

55. M. Collett, How navigational guidance systems are combined in a desert ant. Current Biology 22, 927–932 (2012).

56. S. S. Kim, A. M. Hermundstad, S. Romani, L. F. Abbott, V. Jayaraman, Generation of stable heading representations in diverse visual scenes. Nature 576, 126–131 (2019).

57. Y. E. Fisher, J. Lu, I. D’Alessandro, R. I. Wilson, Sensorimotor experience remaps visual input to a heading-direction network. Nature 2019 576:7785 576, 121–125 (2019).

58. Y. Sun, A. Nern, R. Franconville, H. Dana, E. R. Schreiter, L. L. Looger, K. Svoboda, D. S. Kim, A. M. Hermundstad, V. Jayaraman, Neural signatures of dynamic stimulus selection in Drosophila. Nature Neuroscience 2017 20:8 20, 1104–1113 (2017).

59. M. A. Basnak, A. Kutschireiter, T. S. Okubo, A. Chen, P. Gorelik, J. Drugowitsch, R. I. Wilson, Multimodal cue integration and learning in a neural representation of head direction. Nat. Neurosci. 28, 1729–1740 (2025).

60. C. Dan, B. K. Hulse, R. Kappagantula, V. Jayaraman, A. M. Hermundstad, A neural circuit architecture for rapid learning in goal-directed navigation. Neuron 112, 2581–2599.e23 (2024).

61. P. Mussells Pires, L. Zhang, V. Parache, L. F. Abbott, G. Maimon, Converting an allocentric goal into an egocentric steering signal. Nature 626, 808–818 (2024).

62. J. Ormond, J. O’Keefe, Hippocampal place cells have goal-oriented vector fields during navigation. Nature 2022 607:7920 607, 741–746 (2022).

63. R. G. M. Morris, P. Garrud, J. N. P. Rawlins, J. O’Keefe, Place navigation impaired in rats with hippocampal lesions. Nature 297, 681–683 (1982).

64. J. O’Keefe, J. Dostrovsky, The hippocampus as a spatial map. Preliminary evidence from unit activity in the freely-moving rat. Brain Res. 34, 171–175 (1971).

65. S. H. Singh, F. van Breugel, R. P. N. Rao, B. W. Brunton, Emergent behaviour and neural dynamics in artificial agents tracking odour plumes. Nature Machine Intelligence 2023 5:1 5, 58–70 (2023).

66. S. Vijayabaskaran, S. Cheng, Navigation task and action space drive the emergence of egocentric and allocentric spatial representations. PLoS Comput. Biol. 18, e1010320 (2022).

67. R. A. Andersen, Multimodal integration for the representation of space in the posterior parietal cortex. Philos. Trans. R. Soc. Lond. B Biol. Sci. 352, 1421–1428 (1997).

68. E. Avila, K. J. Lakshminarasimhan, G. C. Deangelis, D. E. Angelaki, Visual and Vestibular Selectivity for Self-Motion in Macaque Posterior Parietal Area 7a. Cereb. Cortex 29, 3932–3947 (2019).

69. E. T. Rolls, Spatial coordinate transforms linking the allocentric hippocampal and egocentric parietal primate brain systems for memory, action in space, and navigation. Hippocampus 30, 332–353 (2020).

70. Y. Kobayashi, D. G. Amaral, Macaque monkey retrosplenial cortex: II. Cortical afferents. Journal of Comparative Neurology 466, 48–79 (2003).

71. T. Hafting, M. Fyhn, S. Molden, M. B. Moser, E. I. Moser, Microstructure of a spatial map in the entorhinal cortex. Nature 2005 436:7052 436, 801–806 (2005).

72. M. S. Madhav, R. P. Jayakumar, B. Y. Li, S. G. Lashkari, K. Wright, F. Savelli, J. J. Knierim, N. J. Cowan, Control and recalibration of path integration in place cells using optic flow. Nature Neuroscience 2024 27:8 27, 1599–1608 (2024).

73. M. S. Maisak, J. Haag, G. Ammer, E. Serbe, M. Meier, A. Leonhardt, T. Schilling, A. Bahl, G. M. Rubin, A. Nern, B. J. Dickson, D. F. Reiff, E. Hopp, A. Borst, A directional tuning map of Drosophila elementary motion detectors. Nature 500, 212–216 (2013).

74. B. Schnell, S. V. Raghu, A. Nern, A. Borst, Columnar cells necessary for motion responses of wide-field visual interneurons in Drosophila. J. Comp. Physiol. A Neuroethol. Sens. Neural Behav. Physiol. 198, 389–395 (2012).

75. K. Shinomiya, A. Nern, I. A. Meinertzhagen, S. M. Plaza, M. B. Reiser, Neuronal circuits integrating visual motion information in Drosophila melanogaster. Curr. Biol. 32, 3529–3544.e2 (2022).

76. M. P. Suver, A. M. M. Matheson, S. Sarkar, M. Damiata, D. Schoppik, K. I. Nagel, Encoding of Wind Direction by Central Neurons in Drosophila. Neuron 102, 828–842.e7 (2019).

77. M. P. Suver, A. M. Medina, K. I. Nagel, Active antennal movements in Drosophila can tune wind encoding. Current Biology 33, 780–789.e4 (2023).

78. A. Kamikouchi, H. K. Inagaki, T. Effertz, O. Hendrich, A. Fiala, M. C. Göpfert, K. Ito, The neural basis of Drosophila gravity-sensing and hearing. Nature 2009 458:7235 458, 165–171 (2009).

79. S. Yorozu, A. Wong, B. J. Fischer, H. Dankert, M. J. Kernan, A. Kamikouchi, K. Ito, D. J. Anderson, Distinct sensory representations of wind and near-field sound in the Drosophila brain. Nature 458, 201–205 (2009).

80. K. E. Cullen, J. S. Taube, Our sense of direction: progress, controversies and challenges. Nature Neuroscience 2017 20:11 20, 1465–1473 (2017).

81. J. L. Fox, A. L. Fairhall, T. L. Daniel, Encoding properties of haltere neurons enable motion feature detection in a biological gyroscope. Proc Natl Acad Sci U S A 107, 3840–3845 (2010).

82. P. Oteiza, I. Odstrcil, G. Lauder, R. Portugues, F. Engert, A novel mechanism for mechanosensory-based rheotaxis in larval zebrafish. Nature 547, 445–448 (2017).

83. G. Valera, D. A. Markov, K. Bijari, O. Randlett, A. Asgharsharghi, J. P. Baudoin, G. A. Ascoli, R. Portugues, H. López-Schier, A neuronal blueprint for directional mechanosensation in larval zebrafish. Current Biology 31, 1463–1475.e6 (2021).

84. K. J. Leitch, F. V. Ponce, W. B. Dickson, F. Van Breugel, M. H. Dickinson, The long-distance flight behavior of Drosophila supports an agent-based model for wind-assisted dispersal in insects. Proc. Natl Acad. Sci. USA 118, e2013342118 (2021).

85. F. Van Breugel, K. Morgansen, M. H. Dickinson, Monocular distance estimation from optic flow during active landing maneuvers. Bioinspir. Biomim. 9 (2014).

86. B. Lingenfelter, A. Nag, F. Van Breugel, Insect inspired vision-based velocity estimation through spatial pooling of optic flow during linear motion. Bioinspir. Biomim. 16, 66004 (2021).

87. A. M. M. Matheson, A. J. Lanz, A. M. Medina, A. M. Licata, T. A. Currier, M. H. Syed, K. I. Nagel, A neural circuit for wind-guided olfactory navigation. Nature Communications 2022 13:1 13, 1–21 (2022).

88. Y. Aso, D. Yamada, D. Bushey, K. L. Hibbard, M. Sammons, H. Otsuna, Y. Shuai, T. Hige, Neural circuit mechanisms for transforming learned olfactory valences into wind-oriented movement. Elife 12, e85756 (2023).

89. A. Hamid, H. Gattuso, A. N. Caglar, M. Pillai, T. Steele, A. Gonzalez, K. Nagel, M. H. Syed, The conserved RNA-binding protein Imp is required for the specification and function of olfactory navigation circuitry in Drosophila. Current Biology 34, 473–488.e6 (2024).

90. N. D. Kathman, A. J. Lanz, J. D. Freed, K. I. Nagel, Neural dynamics for working memory and evidence integration during olfactory navigation in Drosophila. bioRxiv, 2024.10.05.616803 (2024).

91. A. J. Lanz, N. D. Kathman, E. Hao, B. Ermentrout, K. I. Nagel, Disinhibition of a recurrent attractor gates a persistent goal signal for navigation. bioRxiv, 2025.10.07.681003 (2025).

92. A. F. Siliciano, S. Minni, C. Morton, C. K. Dowell, N. B. Eghbali, J. Y. Rhee, L. F. Abbott, V. Ruta, A vector-based strategy for olfactory navigation in Drosophila. bioRxiv, 2025.02.15.638426 (2025).

93. H. Dana, Y. Sun, B. Mohar, B. K. Hulse, A. M. Kerlin, J. P. Hasseman, G. Tsegaye, A. Tsang, A. Wong, R. Patel, J. J. Macklin, Y. Chen, A. Konnerth, V. Jayaraman, L. L. Looger, E. R. Schreiter, K. Svoboda, D. S. Kim, High-performance calcium sensors for imaging activity in neuronal populations and microcompartments. Nature Methods 2019 16:7 16, 649–657 (2019).

94. M. Goslin, M. R. Mine, The Panda3D graphics engine. Computer (Long. Beach. Calif). 37, 112–114 (2004).

95. E. A. Pnevmatikakis, A. Giovannucci, NoRMCorre: an online algorithm for piecewise rigid motion correction of calcium imaging data. J. Neurosci. Methods 291, 83–94 (2017).

96. A. D. Straw, K. Branson, T. R. Neumann, M. H. Dickinson, Multi-camera real-time three-dimensional tracking of multiple flying animals. J. R. Soc. Interface 8, 395–409 (2010).

97. I. Kanitscheider, R. Coen-Cagli, A. Kohn, A. Pouget, Measuring Fisher Information Accurately in Correlated Neural Populations. PLoS Comput. Biol. 11 (2015).

98. C. R. Rao, Information and the Accuracy Attainable in the Estimation of Statistical Parameters. 235–247 (1946).

99. H. Cramér, Mathematical Methods of Statistics (PMS-9). Mathematical Methods of Statistics (PMS-9), doi: 10.1515/9781400883868/HTML (1946).

100. F. Chollet, others, Keras, Github (2015).

